# Cryo-EM structure of the human NKCC1 transporter reveals mechanisms of ion coupling and specificity

**DOI:** 10.1101/2021.11.11.468215

**Authors:** Caroline Neumann, Lena Lindtoft Rosenbæk, Rasmus Kock Flygaard, Michael Habeck, Jesper Lykkegaard Karlsen, Yong Wang, Kresten Lindorff-Larsen, Hans Henrik Gad, Rune Hartmann, Joseph Lyons, Robert A. Fenton, Poul Nissen

**Affiliations:** Danish Research Institute of Translational Neuroscience - DANDRITE, Nordic EMBL Partnership for Molecular Medicine; Department of Molecular Biology and Genetics, Aarhus University, 8000 Aarhus, Denmark; Department of Biomedicine, Aarhus University, 8000 Aarhus, Denmark; Linderstrøm-Lang Centre for Protein Science, Department of Biology, University of Copenhagen, 2200 Copenhagen, Denmark; Shanghai Institute for Advanced Study, Institute of Quantitative Biology, College of Life Sciences, Zhejiang University, Hangzhou 310027, China

**Keywords:** chloride transport, cation:chloride co-transporters, NKCC1, ion coupling, substrate specificity, cryo-EM

## Abstract

The sodium-potassium-chloride transporter NKCC1 (SLC12A2) performs Na^+^-dependent Cl^−^ and K^+^ ion uptake across plasma membranes. NKCC1 is important for regulating e.g. cell volume, hearing, blood pressure, and chloride gradients defining GABAergic and glycinergic signaling in brain. Here, we present a 2.6 Å resolution cryo-electron microscopy (cryo-EM) structure of human NKCC1 in the substrate-loaded (Na^+^, K^+^, 2 Cl^−^) and inward-facing conformation adopting an occluded state that has also been observed for the SLC6 type transporters MhsT and LeuT. Cl^−^ binding at the Cl1 site together with the nearby K^+^ ion provide a crucial bridge between the LeuT-fold scaffold and bundle domains. Cl^−^ ion binding at the Cl2 site seems to undertake a structural role similar to a conserved glutamate of SLC6 transporters and may allow for chloride-sensitive regulation of transport. Supported by functional studies in mammalian cells and computational simulations we describe the Na^+^ binding site and a putative Na^+^ release pathway along transmembrane helix 5. The results provide insight into the structure-function relationship of NKCC1 with broader implications for other SLC12 family members.

## Introduction

Eukaryotic cation:chloride co-transporters (CCCs) belonging to the SLC12 family can be divided into three subclasses: the sodium:potassium:chloride (NKCCs, SLC12A1-2), sodium:chloride (NCC, SLC12A3) and potassium:chloride (KCCs, SLC12A4-7) co-transporters. Furthermore, there are two orphan transporters CCC9 (SLC12A8) and CIP1 (SLC12A9) (Arroyo *et al*, 2013). The Cl^−^ translocation by CCCs is secondary active, driven by Na^+^ and/or K^+^ gradients established by the plasma membrane Na^+^-K^+^-ATPase (Markadieu & Delpire, 2014; Payne, 2012). NKCCs transport K^+^ and Cl^−^ into cells coupled to Na^+^ influx, NCC transports Na^+^ and Cl^−^ into cells, whereas KCCs extrude K^+^ and Cl^−^ out of cells (Hebert *et al*, 2004). All the characterized co-transporters are electroneutral and are important for maintaining cellular Cl^−^ balance and cell volume in numerous different cell types (Delpire & Gagnon, 2018). NCC and NKCC2 activity in the kidney are important for maintenance of NaCl homeostasis and hence blood pressure regulation (Arroyo *et al.*, 2013), with NKCC2 also important for NH_4_^+^ reabsorption and acid-base homeostasis (Markadieu & Delpire, 2014). Maintenance of cellular Cl^−^ balance by CCCs in the central nervous system is not only important for neuronal proliferation and differentiation, but it determines the strength of inhibitory, hyperpolarizing neurotransmission by glycine- and GABA-gated chloride channels, respectively (Blaesse *et al*, 2009) or even reverse polarities. NKCC1 also inhibits the leucine transporter LAT1 and Akt/Erk pathways and mTORC1 activation (Demian *et al*, 2019), thereby creating a connection between cell volume and cell mass regulation. The importance of CCCs is highlighted by numerous diseases, with inactivating mutations of NKCC2 leading to type I Bartter’s syndrome (Simon *et al*, 1996a), mutations of NCC causing Gitelman syndrome (Ji *et al*, 2008; Simon *et al*, 1996b), both characterized by reduced K^+^ levels in blood and elevated blood pH. Mutations of KCC3 cause Andermann syndrome, a severe neurodegenerative disorder (Arroyo *et al.*, 2013). NKCC1 mutations are also linked to impaired hearing and neurodevelopment (Koumangoye *et al*, 2021). Furthermore, improper functioning of CCCs are associated with various neurological and psychiatric disorders, such as epilepsy, neuropathic pain, anxiety, cerebral ischemia, autism and schizophrenia (Jaggi *et al*, 2015).

Recently, cryo-EM structures of the dimeric NKCC1 (Chew *et al*, 2019; Yang *et al*, 2020; Zhang *et al*, 2021), KCC1 (Liu *et al*, 2019), KCC2 (Chi *et al*, 2021; Xie *et al*, 2020), KCC3 (Xie *et al.*, 2020) (Chi *et al*, 2020) and KCC4 (Xie *et al.*, 2020) transporters as well as a monomeric structure of KCC4 (Reid *et al*, 2020) have described the three-dimensional architecture of these transporters, their fold and the potential localization of their Na^+^ and/or Cl^−^ binding sites. All these structures present the transporters in their partially occluded inward-facing conformation with substrates still occupying the binding pockets. They contain 12 transmembrane (TM) helices with an inverted pseudo-twofold symmetry between TM1-5 and TM6-10, first identified as the LeuT-fold (Yamashita *et al*, 2005). TM helices 1 and 6 contain non-helical junctions in the middle that allow for independent movement of the intracellular and extracellular halves to expose or close the substrate-binding sites to solvent access in an ‘alternating access’ manner (Joseph *et al*, 2019). Despite coming from two distinct vertebrates with only 74.2% sequence identity, the K^+^ and Cl^−^ binding sites in zebrafish (*Danio rerio*, zNKCC1) and human NKCC1 (hNKCC1) are fully conserved, highlighting the importance of specific residues for interacting with the transported ions (Chew *et al.*, 2019; Yang *et al.*, 2020). The regulatory N-terminal domain in hNKCC1 is 77 amino acid residues longer than in zNKCC1 (285 vs 208 residues) and is disordered in nature^12^. Crystal structures of the regulatory C-terminal domain of a bacterial NKCC1 homologue (Warmuth *et al*, 2009) and of the eukaryotic C-terminal domain of KCC1 have provided further insight into the regulatory mechanisms of CCC transporters (Zimanyi *et al*, 2020).

Structural studies of amino acid-polyamine-organocation (APC) transporters adopting the LeuT fold have provided extensive insight and understanding of their transport mechanisms, substrate specificity, ion coupling and inhibitory action. Since the first LeuT crystal structure (Yamashita *et al.*, 2005), several other SLC6 structures (MhsT (Malinauskaite *et al*, 2014), dDAT (Penmatsa *et al*, 2013), SERT (Coleman *et al*, 2016), B0AT1 (Yan *et al*, 2020), GlyT (Shahsavar *et al*, 2021)) have been determined in a variety of different conformational states and in complex with substrates or inhibitors. Additionally, structures of sequence-unrelated, but LeuT fold adopting SLC families have also been described, e.g. vSGLT (SLC5) (Faham *et al*, 2008; Watanabe *et al*, 2010), GkApcT (SLC7) (Jungnickel *et al*, 2018), ScaDMT (SLC11) (Ehrnstorfer *et al*, 2014), NKCC1 (SLC12) (Chew *et al.*, 2019; Yang *et al.*, 2020; Zhang *et al.*, 2021) and SiaT (SLC35) (Wahlgren *et al*, 2018). Interestingly, despite minimal sequence similarity between different APC transport families, mechanistic aspects of Na^+^ dependent transport seem to be retained among many of them. All the transporters contain two domains: the bundle domain comprising TM 1-2 and TM 6-7 that performs the main conformational changes throughout the transport cycle, and the scaffold domain encompassing TM 3-4 and TM 8-9 that stays almost rigid throughout the functional cycle. This is often referred to as a rocking bundle mechanism (Forrest *et al*, 2011; Forrest & Rudnick, 2009). Ligand and ion binding sites are localized at the interface of the scaffold and bundle domain and binding/release of specific ligands control bundle movements and coupling (Zhang *et al*, 2018). For the well-characterized SLC6 transporters, binding of extracellular Na^+^ at the Na2 site (which also corresponds to the expected Na^+^ binding site of NKCC1) stabilizes outward-facing states, followed by subsequent binding of substrates at S1 and the Na1 site. The Na^+^ and substrate bound state can then explore conformational changes towards the inward-facing state (Ben-Yona & Kanner, 2009; Claxton *et al*, 2010; Fenollar-Ferrer *et al*, 2014; Kazmier *et al*, 2014; Tavoulari *et al*, 2016; Zhao *et al*, 2010), where Na^+^ release from the Na2 site to the cytoplasm (Malinauskaite *et al.*, 2014) leads further to the inward-open state, where substrates are released from the S1 and Na1 sites (Krishnamurthy & Gouaux, 2012). The main substrate alone has no effect on the conformational state without sodium ions present, and the Na2 site is a crucial structural element for cytoplasmic occlusion and opening (Tavoulari *et al.*, 2016).

Here, we present a cryo-EM structure of the full-length human NKCC1 transporter in an occluded, inward-oriented state in complex with K^+^, two Cl^−^ ions and Na^+^ (Fig. S1A-B). Using the high-resolution maps coupled to functional analysis in mammalian cells we characterize the ion and solvent binding sites of NKCC1, and further explore substrate specificity, coupling and release of ions in NKCC1 and their relationship to other CCCs. We also investigate the potential function of the Na^+^ release pathway leading to the cytoplasmic environment and make comparisons of the NKCC1 structure with SLC6 type transporters matching similar structural states and sites.

## Results

### NKCC1 cryo-EM determination

We collected cryo-EM data of hNKCC1, which combined with single-particle analysis, resulted in a 2.6 Å reconstruction of the transmembrane domain of human NKCC1 in its occluded, inward-facing, substrate-bound conformation. i.e. a state preceding release of substrates to the cytoplasm (Fig. 1A). In three individual data sets, single particle images were extensively cleaned through multiple rounds of 3D classification before high-resolution reconstructions were obtained from 3D auto-refinement of Bayesian polished particles. Inspection of the reconstructions from the individual data sets revealed the same state of hNKCC1, thus we merged data into a single stack of 256,791 particle images to obtain a higher-quality reconstruction (Fig. EV1, EV2 and EV3). Similar to other NKCC1 cryo-EM structures (Chew *et al.*, 2019; Yang *et al.*, 2020; Zhang *et al.*, 2021), the N-terminal domain could not be identified in the map. The regulatory cytoplasmic domain (aa 756-1212) appears to be flexible and despite extensive attempts for classification in cryoSPARC (Punjani *et al*, 2017) and the use of multibody refinement in RELION (Nakane *et al*, 2018), it appeared only at low resolution locally (7-8Å). However, it confirmed an overall asymmetric configuration of the transmembrane domains (Fig. 1A and EV3C). The cytoplasmic domains were masked out along with the detergent micelle and the refinement was focused on the transmembrane part of the transporter (Fig. EV1 and EV2).

**Fig. 1.**
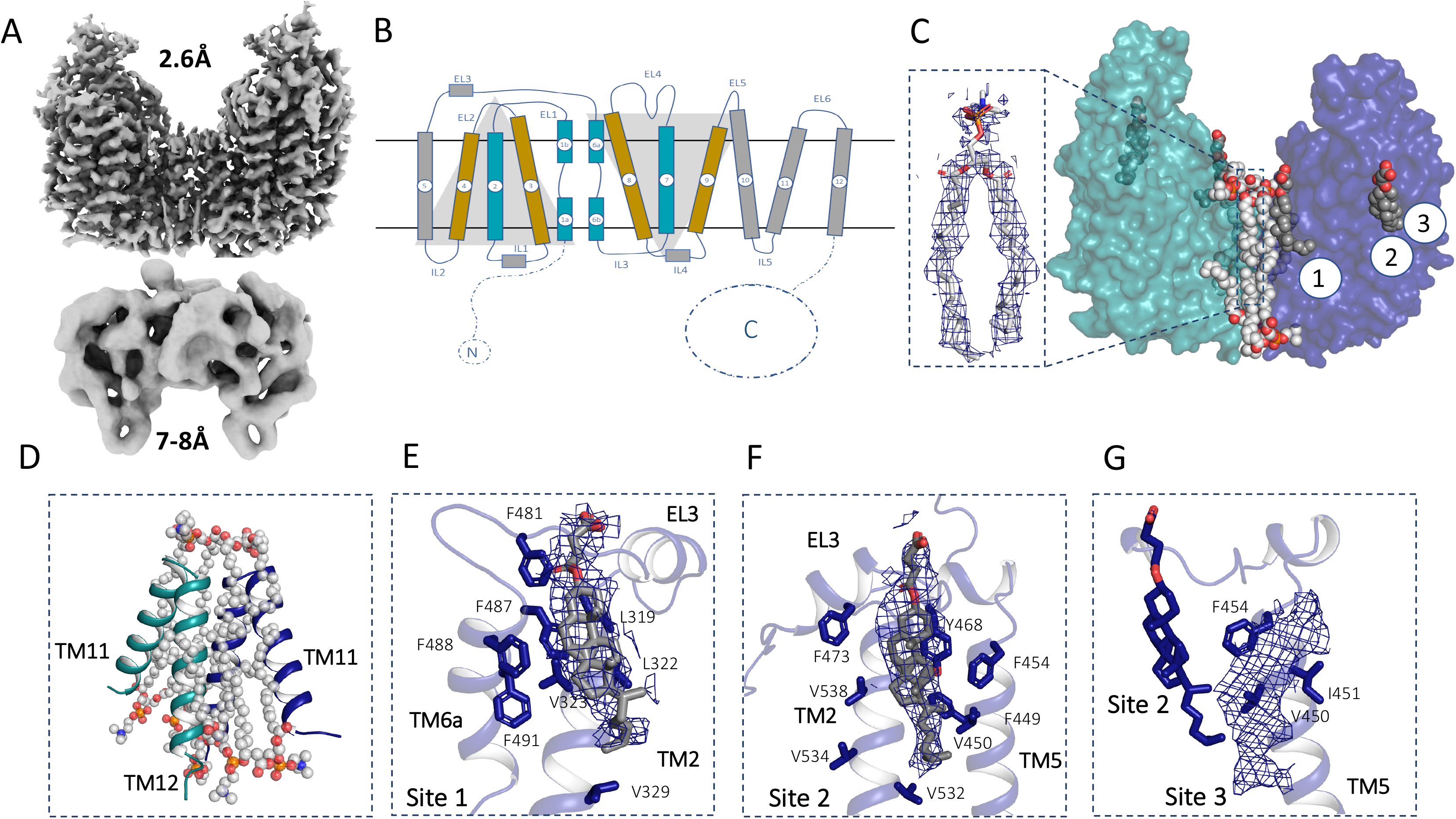
Overview of hNKCC1 structure. A) EM density of the transmembrane domain of hNKCC1 (resolved to 2.6Å) and the cytoplasmic domain (resolved to 7-8Å). B) Schematic representation of the NKCC1 structure (LeuT fold). Scaffold helices are presented in wheat, bundle helices in cyan and the remaining helices presented in grey. C) Surface representation of the transmembrane domain of NKCC1 with lipid molecules localized at the dimerization interface (shown in spheres) and the three cholesterol binding sites (shown in spheres). In the inset, a representative density for a lipid molecule is presented. D) TM11 and TM12 from each protomer are presented together with the lipid molecules present at the dimerization interface. E, F, G) Densities for cholesterol binding site 1, site 2 and potential site 3 are presented together with the modelled cholesterol hemisuccinate molecules.

The resolution of the final map was high enough to clearly identify non-protein features corresponding to a K^+^ ion and two Cl^−^ ions. Density at the Na^+^ binding site appears at a lower, but significant contour level, indicating that the Na^+^ site is also occupied and saturated at the 200 mM Na^+^ concentration of the purification buffer. When compared to the well-characterized SLC6 type amino acid transporters LeuT (Gotfryd *et al*, 2020) and MhsT (Malinauskaite *et al.*, 2014), it is clear that despite different substrates the substrate binding sites are similar and hence the mechanisms of Na^+^ coupled transport appear mutually informative.

### Lipid and cholesterol binding to hNKCC1

The overall predicted LeuT-fold topology and dimeric architecture of the transmembrane region of the determined hNKCC1 are presented in Fig. 1B-C. Similar to zNKCC1 (Chew *et al.*, 2019), our three-dimensional EM map revealed the presence of seven lipid molecules bound at the central cleft with either one or both acyl chains inserted between the two protomers (Fig. 1C-D). One lipid is placed directly on the C2 symmetry axis, whereas the remaining lipid molecules cluster in two pairs of 3 annular lipids found on opposite sides of the interface.

The EM map identified three cholesterol binding sites per NKCC1 protomer at the transporter surface, corresponding to the outer leaflet of the plasma membrane (Fig. 1C). Therefore, we decided to model cholesteryl hemisuccinate molecules (CHS, used in the solubilization and purification) into the densities. The first CHS molecule was found close to the dimer interface, binding at a groove formed between TM2 and TM6a with its *α*-face forming a *π*-*π* interaction with F487 on TM6a. The CHS molecule is also accommodated by aromatic residues on TM6a (F488 and F491), in addition to F481 on EL3 and non-polar residues (L319, L322, V323, V329) on TM2 (Fig. 1E). The second CHS molecule interacts with hydrophobic residues on TM2 (V532, V534 and V538), TM5 (F449, V450 and F454) and F473 on EL3 and it forms a *π*- *π* interaction with Y468 on EL3 (Fig. 1F). A third potential CHS molecule was identified in close proximity, interacting hydrophobically with the second CHS molecule, the non-polar V450 and I451 on TM5 and forming a *π*- *π* interaction with F454 on TM5 (Fig. 1G). The molecule was, however, not modelled into the map. Assuming that hNKCC1 translocates its substrates using the rocking bundle mechanism similarly to other transporters adopting the LeuT-fold, the three CHS molecules appear to stabilize the inward-facing conformation of the transporter by both blocking movement of the extracellular parts of the bundle domain (TM2, TM5, TM6a) and disfavoring transitions to the outward-facing conformation with an open extracellular vestibule. Comparison of the unmodelled densities from the deposited KCC3 cryo-EM structure (Chi *et al.*, 2021) reveal that all three cholesterol binding sites are conserved, despite different amino acid composition within these sites (Fig. S1D and E). A potential fourth cholesterol binding site could also be identified in KCC3 (Fig. S1F), where a cholesterol molecule can be accommodated in a groove formed by TM5 and TM7. Therefore, it seems that cholesterol might be of importance for both sodium-dependent and sodium-independent CCC transporters, stabilizing the substrate releasing state of NKCC1 and the substrate loading state of KCC transporters. These observations also support the association of both NKCC1 and KCC2 with cholesterol rich lipid rafts in the plasma membrane, where cholesterol could regulate their transport cycle (Hartmann *et al*, 2009).

### Sodium ion binding site and its role in ion coupling

In hNKCC1, the Na^+^ ion is coordinated by the main chain carbonyl oxygen of W300 and L297 on TM1 of the bundle domain, and the side chain hydroxyl groups of S613 and S614 and the main chain carbonyl oxygen of A610 on TM8 of the scaffold domain (Fig. 2A). This corresponds to the Na2 driver site of SLC6 type transporters like LeuT, dDAT, GlyT1, SERT and MhsT (Yamashita *et al.*, 2005) (Fig. 2C). Contrasting the previously described zNKCC1 and hNKCC1 structures (Chew *et al.*, 2019; Yang *et al.*, 2020; Zhang *et al.*, 2021), we consider the Na^+^ bound hNKCC1 as an occluded inward-oriented conformation, which represents a trapped intermediate preceding Na^+^ and subsequently K^+^ and Cl^−^ release to the cytoplasmic environment. When comparing our hNKCC1 to the SLC6-type transporters MhsT and LeuT in forms that also adopt this inward-oriented and occluded state, we find striking similarities (Fig. 2C and Fig. S2). Most notably, we observe a clear candidate release pathway for Na^+^ that extends from the Na^+^ binding site to the cytoplasm along TM5. This pathway represents the critical, first encounter of the Na^+^ site with the cytoplasmic environment, leading to subsequent solvation and release of Na^+^ into the low-sodium cytoplasmic environment, hence defining the vectorial flow of transport. Defining the dynamics and functional properties of this pathway would provide key characteristics of NKCC1 function.

**Fig. 2.**
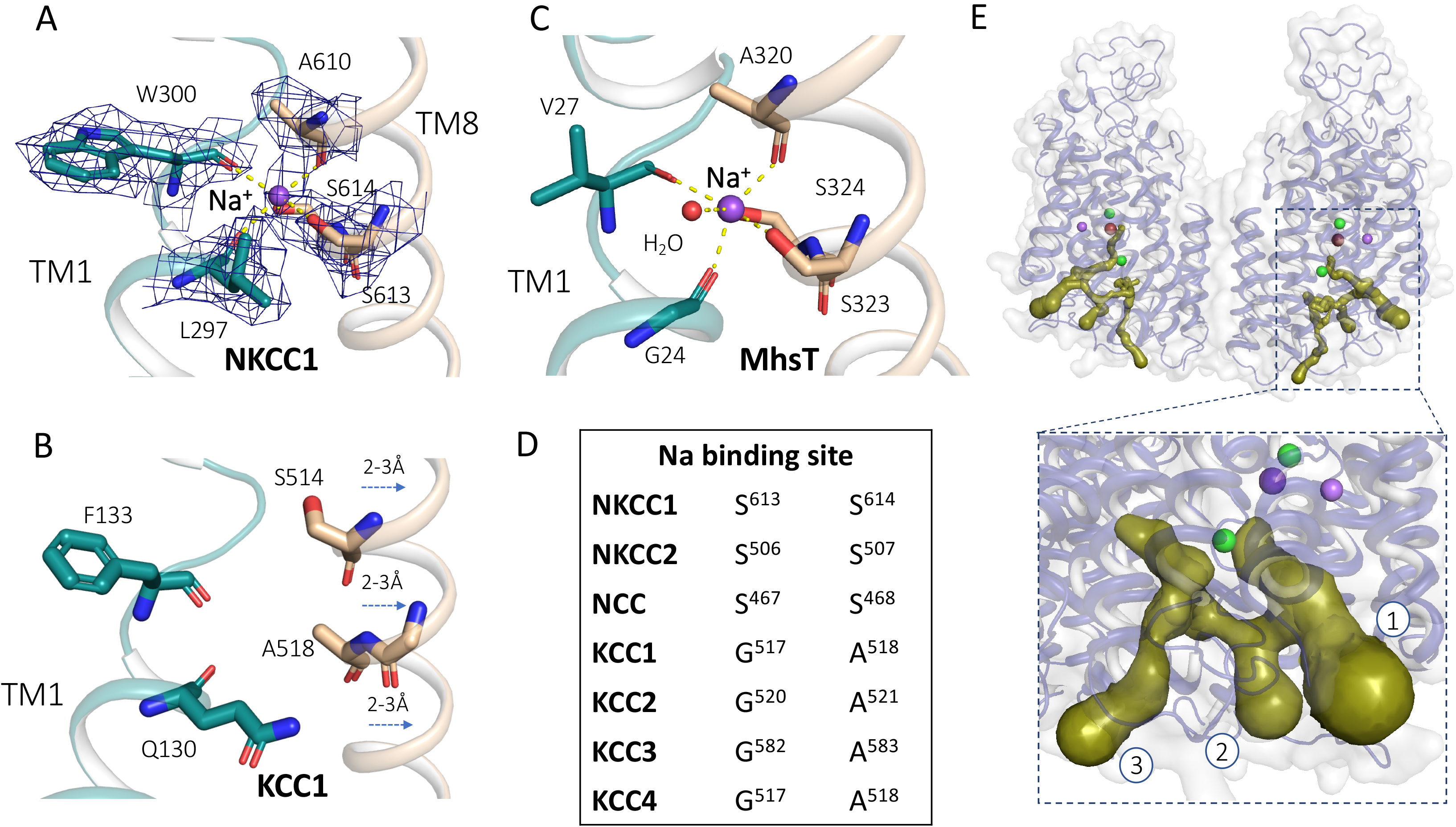
Sodium binding site. A) The sodium binding site in hNKCC1. The ion has trigonal-bipyramidal coordination by five ligands. Scaffold helices are presented in wheat and bundle helices are presented in cyan; the sodium ion is presented as a purple sphere and coordinating residues are presented as sticks. B) KCC1 does not bind sodium due to a G-A substitution on TM8. Additionally, the helix is moved 2-3Å away when compared with hNKCC1. C) The sodium binding site in MhsT. The ion is solvated and therefore, it has distorted octahedral coordination by five amino acid residues on TM1 and TM8 and a water molecule acting as the sixth ligand. D) Sequence alignment of the sodium site of the CCC transporters. The G-A substitution of KCC1-4 on TM8 is responsible for abolishment of sodium binding. E) Three potential intracellular exit pathways predicted by MOLE 2.5.

Water molecules are found within the intracellular vestibule, but in contrast to the occluded, inward-facing structure of MhsT (Malinauskaite *et al.*, 2014), our hNKCC1 structure shows no solvation of the Na^+^ site, like the occluded LeuT structures (Malinauskaite *et al*, 2016; Yamashita *et al.*, 2005) (Fig. 2 and Fig. S3). The MOLE2.5 program was used to identify and analyze potential intracellular exit pathways (Berka *et al*, 2012) (Fig. 2E), as also described previously for hNKCC1^K289N_G351R^ (Yang *et al.*, 2020). Interestingly, intracellular pathway 1 is lined by E429 and E431 (predicted pKa-values of 4.6 and 4.5 by PROPKA, i.e. both negatively charged at pH 7.5) at the cytoplasmic interface (Fig. 3A). These residues are conserved in hNKCC1, hNKCC2 and hNCC, but absent in the sodium-independent KCC transporters (Fig. 3C). Thus, they may act as internal “gatekeepers” that attract the Na^+^ ion from the binding site into the cytoplasmic environment, in a similar manner to as observed in cation channels (Doyle *et al*, 1998). Bacterial, but not mammalian, SLC6 transporters contain a conserved glutamate residue at a similar position to E429 (E192^LeuT^) that interacts with and escorts the Na^+^ out of the transporters (Shaikh & Tajkhorshid, 2010) (Fig. 3B). This residue also corresponds to the conserved D182 in vSGLT (SLC5) that also facilitates Na^+^ diffusion (Li & Tajkhorshid, 2009). To probe further a putative role of E429 and E431 in Na^+^ release for hNKCC1, Tl^+^ influx experiments (mimicking K^+^ uptake) were performed in mammalian cells expressing wild-type hNKCC1 or various hNKCC1 mutants. The relative rate of Tl^+^ influx was significantly lower for E429A, E431A and E431Q mutants when compared to wild-type hNKCC1. Additionally, there was no statistically significant difference between the relative transport rates of E431A (mutation to a non-polar residue without a negative charge) and E431Q mutants (mutation to a polar residue without a negative charge) (Fig. 3D). Together these results suggest important roles of E429 and E431 in hNKCC1 activity. Cell-surface expression of wild-type hNKCC1 and hNKCC1 mutants were not significantly different, indicating that the altered activity observed with the mutants is not due to altered association of the mutant protein with the plasma membrane (Fig. EV4C-D).

**Fig. 3.**
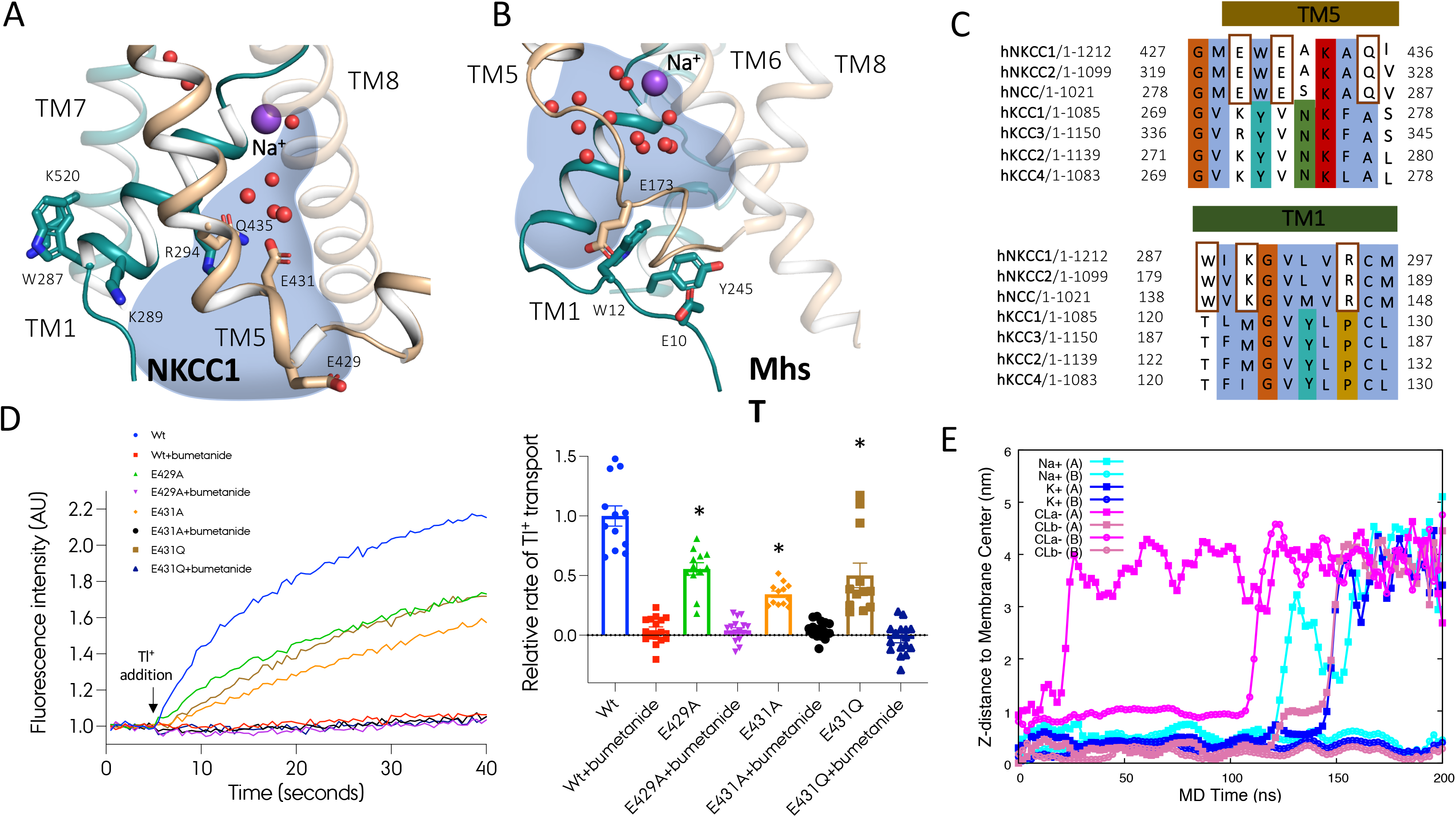
Intracellular sodium release pathway. A) The intracellular sodium release pathway of hNKCC1. The pathway is filled with water-molecules (red spheres) and lined with two negatively charged glutamate residues (Glu429 and Glu431). B) The intracellular sodium release pathway in MhsT filled with water molecules that solvate the sodium ion. C) Sequence alignment of the intracellular part of TM5 and of TM1a of CCC transporters. The sodium-dependent CCC transporters contain two conserved glutamate and a glutamine residue within TM5 and a positively charged lysine and arginine residues on TM1a. D) Transport kinetics and relative initial transport rates of wild-type hNKCC1 and NKCC1 with a mutation within the potential sodium release pathway (E429A, E431A, E431Q) with and without addition of bumetanide; mean ± SEM, n= 3 independent experiments. In example of individual traces, fluorescence intensity is normalized to the average initial baseline period (without Tl^+^) for each individual cell line. In summary data, transport rates are normalized to individual NKCC1 expression per cell line and subsequently normalized to wild-type NKCC1 without bumetanide. E) Substrate release order determined by MD simulations. The Cl^−^ bound at Cl2 leaves first, then the Na^+^ ion, followed by K^+^ and Cl^−^ from the main binding site.

The hNKCC1 structure revealed interactions between R294 on TM1a and Q435 on TM5 (Fig. 3A). Similar interactions are found in SLC6 structures with a sealed intracellular side, corresponding to the outward-oriented states of LeuT (Yamashita *et al.*, 2005) and the occluded inward-facing structure of MhsT (Fig.3 and Fig. S3) (Malinauskaite *et al.*, 2014). The arginine and glutamine residues in hNKCC1 are exclusively conserved in sodium-dependent CCCs suggesting that they are important for Na^+^ transport, with potentially a similar function to SLC6 transporters (Fig. 3C). Consistent with this model, a R294A mutant has reduced transport activity relative to wild-type NKCC1 (Yang *et al.*, 2020). K289 on TM1 is also conserved in the sodium-dependent CCCs, providing a potential interaction partner for one of the two conserved glutamate residues on TM5 in outward-facing conformational states, where the intracellular side is fully sealed.

Sequence alignment of the CCCs reveals no helix-breaking motif in the intracellular half of TM5, suggesting that TM5 stays intact in the SLC12 family, unlike for SLC6 transporters (Malinauskaite *et al.*, 2014) (Fig. S4). This indicates that NKCC1, NKCC2 and NCC have different dynamics for Na^+^ site solvation and Na^+^ release, potentially involving a more pronounced movement of TM1a (Fig. S3).

### Coupling network within the main substrate binding pocket

The substrate site in hNKCC1 accommodates a K^+^ and Cl^−^ ion, as previously demonstrated for the zNKCC1 (Chew *et al.*, 2019), hNKCC1 (Zhang *et al.*, 2021), hKCC1 (Liu *et al.*, 2019), hKCC2 (Chi *et al.*, 2020), hKCC3 (Chi *et al.*, 2020) and hKCC4 (Reid *et al.*, 2020; Xie *et al.*, 2020) by coordination chemistry and molecular dynamics (MD) simulations. The K^+^ ion is coordinated by the carbonyl oxygens of N298, I299, T499, P496 and the hydroxyl groups of Y383 and T499 (Fig. 4A). Y383 seems to have an important role in substrate specificity, as relative to NKCCs and KCCs, the conserved site in NCC (not transporting K^+^) is a histidine residue, which may substitute for K^+^ (Fig. 4C) (Hartmann & Nothwang, 2014). Previous functional studies demonstrated that mutation of the tyrosine residue results in impairment of transport of the human NKCC1 (Somasekharan *et al*, 2012). The corresponding Y108^LeuT^ and Y176^SERT^ are crucial residues for substrate induced conformational change from the outward-facing to the inward-facing states in SLC6 transporters and participate in an interaction network between the bundle (TM 6 and 7) and scaffold domain (TM 3) in the extracellular pathway (Zhang *et al.*, 2018) (Fig. 4B). In hNKCC1, this network involves Y383 on TM3 that coordinates the K^+^ ion as well as T499 on the unwound part of TM6. Highlighting the importance of these residues, Tl^+^ influx in mammalian cells expressing hNKCC1 Y383F and Y383S mutants was impaired when compared to wild-type hNKCC1 (Fig. 4D), despite similar cell-surface expression of the mutants and wild-type NKCC1 (Fig. EV4 C-D).

**Fig. 4.**
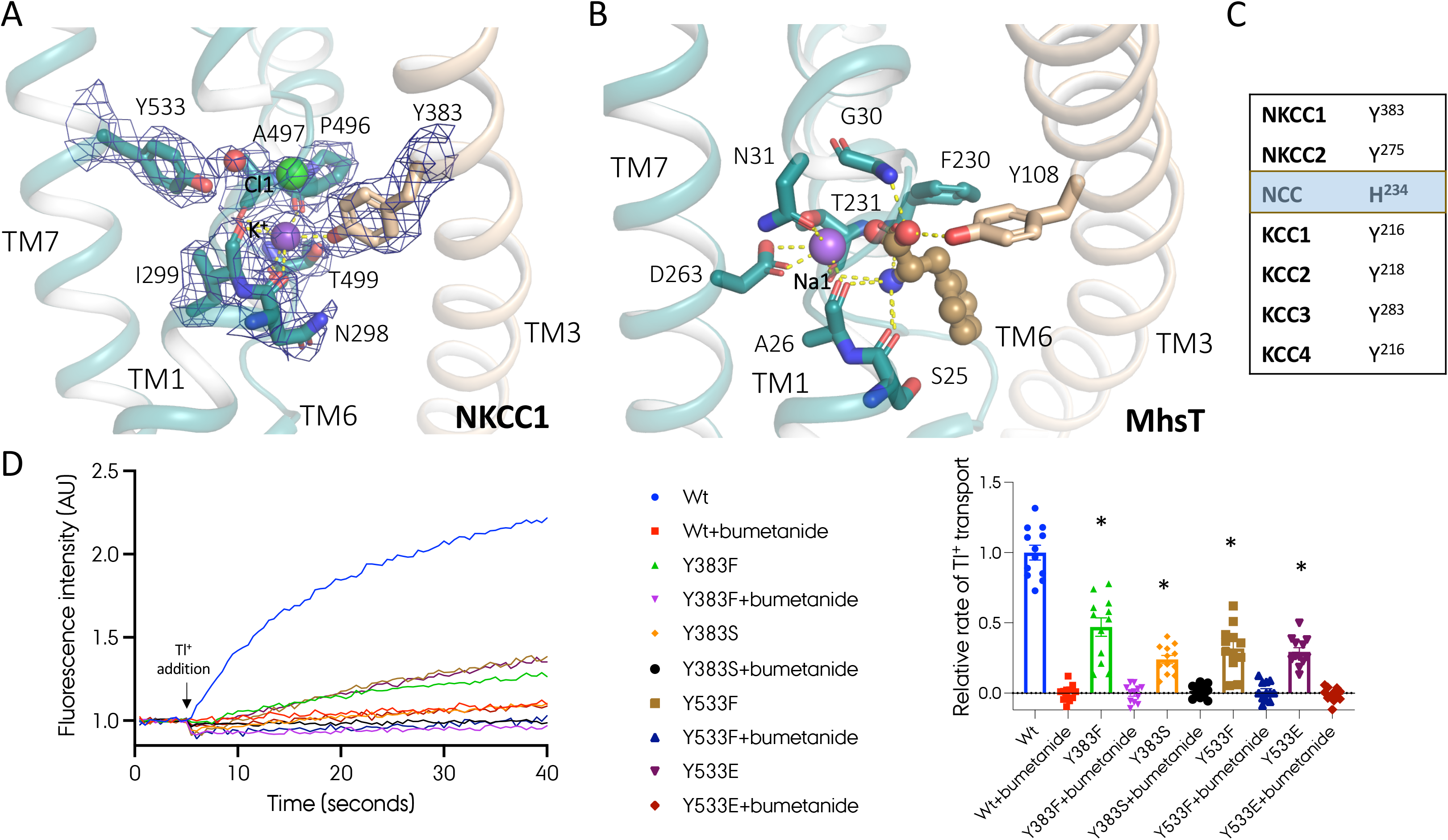
Coordination network within the main binding pocket characterized for. A) The main binding site of hNKCC. The binding pocket accommodates a potassium and chloride ion coordinated by TM3 from the scaffold domain (wheat) and TM1, TM6 and TM7 from the bundle domain (cyan). The density of the ions and residues important for substrate coordination is presented. B) MhsT in complex with L-tryptophan and two sodium ions. Even though the transporter translocates significantly different substrates than the SLC12 transporters, the coordination network remains similar. The substrates bridge interactions between TM3 from the scaffold domain (wheat) and TM1, TM6 and TM7 from the bundle domain (cyan). C) Sequence alignment of the crucial amino acid residue on TM3 of CCC transporters. In contrast to other members of the SLC12 family that contain a tyrosine residue coordinating the K^+^ ion at this position, hNCC contains a histidine residue that in its charged state could substitute the potassium ion. D) Transport kinetics and relative initial transport rates of wild-type hNKCC1 and NKCC1 with a mutation within the K- and Cl1-binding pocket (Y383F, Y383S, Y533F and Y533E) with and without addition of bumetanide; mean ± SEM, n= 3 independent experiments. See Fig 3 legend for normalization approaches.

On the opposite side of the binding pocket, Y533 on the bundle domain of TM7 interacts with A497 on TM6 as well as the Cl^−^ ion through a water-mediated interaction (Fig. 4A). A substrate site coordination network has previously been described for the SLC6 transport family (Zhang *et al.*, 2018). However, in contrast to the Y533 on TM7 in hNKCC1, the prokaryotic SLC6 homologues contain a conserved E290^LeuT^ on TM7 and eukaryotic SLC6 transporters contain a serine residue on TM7 that interacts with TM6 through a chloride ion (Coleman *et al.*, 2016; Penmatsa *et al.*, 2013). TM6 of SLC6 transporters interacts with the Na^+^ ion at Na1 and the amine group of the amino acid substrate. Therefore, it seems that Y533 of hNKCC1 undertakes a role that is similar to both E290^LeuT^ (or a Cl^−^ site) and the Na1 sodium ion in SLC6 transporters. Supporting that this tyrosine residue is crucial for proper substrate translocation by NKCC1, replacing Y533 in hNKCC1 with a phenylalanine or glutamate residue abolishes Tl^+^ influx in transfected mammalian cells (Fig. 4D).

### Second chloride binding site (Cl2) is solvated

Besides the Cl^−^ ion bound at the main substrate site, NKCC1 transports a second Cl^−^ ion accommodated at the Cl2 binding site further towards the cytoplasmic interface. The Cl^−^ ion is coordinated by Y686 on TM10 and additionally, it interacts with the unwound part of TM6 (Fig. 5A). In contrast to the conservation of the Na2 binding site throughout most APC transporters, the two Cl^−^ sites present in hNKCC1 are not found in any other APC transporters. Interestingly, the Cl2 site of hNKCC1 superimposes with the carboxylate side chain group of a strictly conserved glutamate residue on TM2 of SLC6 transporters (e.g. E136^SERT^ and E62^LeuT^), which is important for substrate transport and conformational changes (Chen *et al*, 2001; Korkhov *et al*, 2006; Sen *et al*, 2005; Sucic *et al*, 2002) (LeuT shown in Fig. 5B). Replacement of Y686 with a serine or glutamate residue substantially reduces transport activity of NKCC1 indicating that the residue is crucial for coordination of the chloride ion and maintaining the coupling network between the main binding pocket and distal parts of the transporter (Fig 5E). In hNKCC1, the Cl^−^ ion bridges an interaction between TM10 and TM6, whereas the conserved glutamate residue of SLC6 transporters connects TM2 with TM6 and interacts with another glutamate on TM10 (e.g. E508^SERT^ and E419^LeuT^). An interaction between TM2 and TM10 is also present in CCC transporters as N690 in hNKCC1 on TM10 interacts with two residues on TM2 (S337 and T334) (Fig. 5A and C). Importantly, and in contrast to the unsolvated sodium ion, the Cl^−^ ion bound at Cl2 interacts with cytoplasmic water molecules (Fig. 5A).

**Fig. 5.**
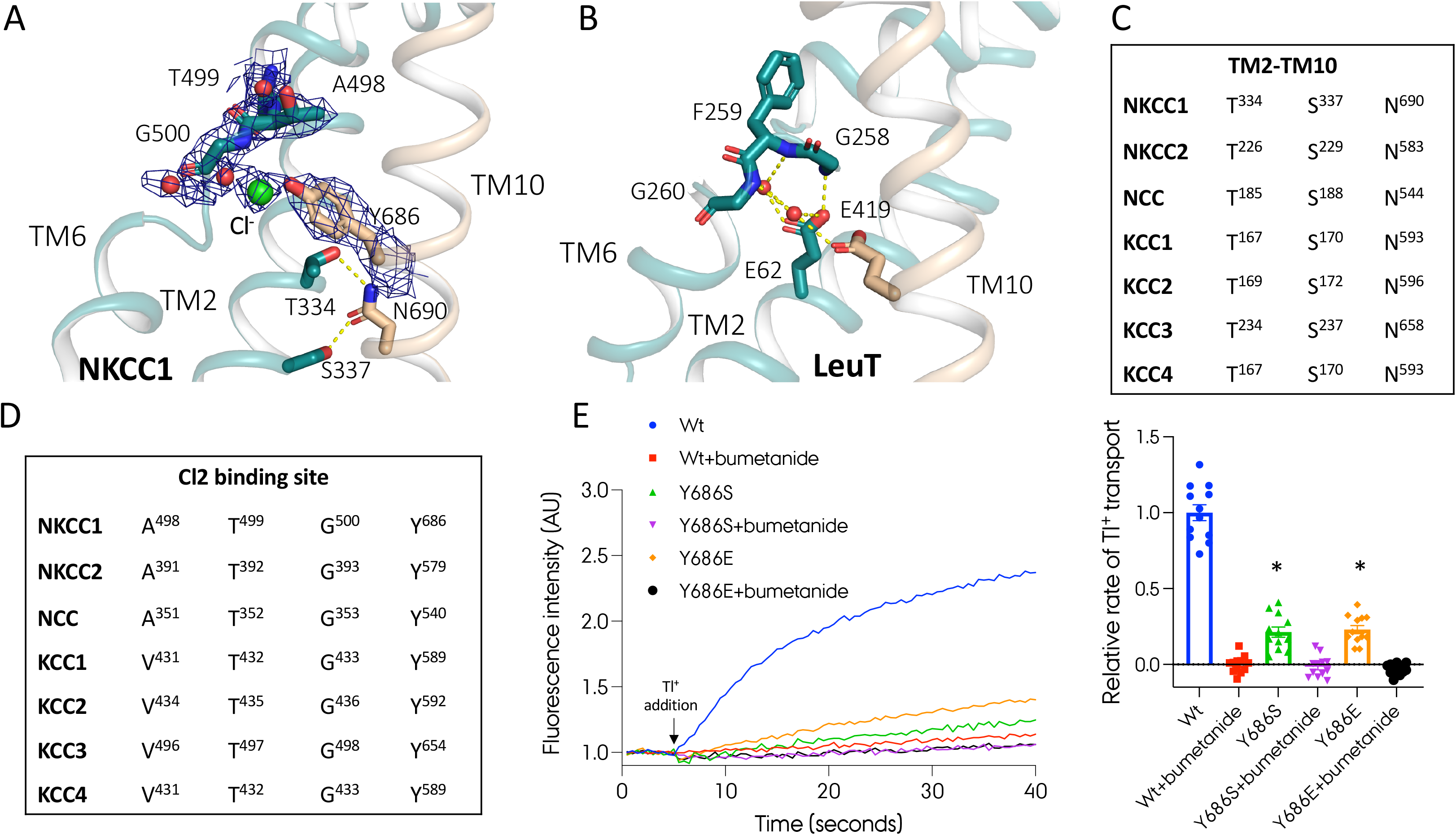
Coordination network between TM2-TM10 and the unwound part of TM6 in. A) Cl2 binding site with the density map. The chloride ion is coordinated by the scaffold helix TM10 (wheat color) and the unwound part of TM6 belonging to the bundle domain (cyan color). B) LeuT in complex with L-leucine and sodium C, D) Sequence alignment of residues linking TM2 and TM10 and residues within the Cl2 binding site. E) Transport kinetics and relative initial transport rates of wild-type hNKCC1 and NKCC1 with a mutation within the Cl2 site (E429A, E431A, E431Q) with and without addition of bumetanide; mean ± SEM, n= 3 independent experiments. See Fig 3 legend for normalization approaches.

### Substrate release

The well-defined ion binding sites with solvation of the Cl2 site prompted us to investigate the substrate-bound complex by MD simulations. We performed twenty independent all-atom MD simulations of the transmembrane domain of our dimeric hNKCC1 embedded into a mixed lipid bilayer consisting of phosphatidylcholine (POPC)/phosphoethanolamine (POPE)/cholesterol at a ratio of 2:2:1, resulting in 8 μs accumulated trajectories. In agreement with observation of hydration of Cl^−^ at the Cl2 site, the solvated Cl^−^ ion was not stably bound and was able to leave its binding site for tens of ns and exchange for solvent Cl^−^ ions. However, in contrast to the reversible release of Cl^−^, a subsequent release of the Na^+^ ion after release of the Cl^−^ ion from Cl2 marks a point of no return, and neither the Cl^−^ or the Na^+^ ions will rebind to their respective sites thereafter. Additionally, in some of our simulations we also observed that the K^+^ substrate and the Cl^−^ ion at Cl1 site left the binding sites almost at the same time, although complete substrate release events in the hNKCC1 dimer were rarely observed probably due to limited simulation time (200 ns for each) (Fig. 3E). Our atomistic MD simulations support a substrate release order in which Cl^−^ at Cl2 site first releases, then Na^+^, and lastly Cl^−^ at the Cl1 site and K^+^ in a cooperative manner.

## Discussion

Structures of NKCC1 (Chew *et al.*, 2019; Yang *et al.*, 2020; Zhang *et al.*, 2021) and KCC1-4 (Chi *et al.*, 2021; Liu *et al.*, 2019; Reid *et al.*, 2020; Xie *et al.*, 2020) indicate that the SLC12 family belongs to the APC superfamily with a LeuT-fold architecture (Fig. 1A). Here, we report a higher resolution structure of human NKCC1 in the substrate-bound form adopting an inward-oriented and occluded state. The Na^+^ ion of NKCC1 binds at the conserved Na2 site, the K^+^ and Cl^−^ ion (Cl1) bind at the general substrate binding pocket of the APC superfamily and the second Cl^−^ ion (Cl2) is found at the same location as a conserved glutamate residue of SLC6 transporters.

Our cryo-EM maps revealed the presence of interfacial lipids, which are also found in the zNKCC1 structure (Chew *et al.*, 2019). These interfacial lipid molecules appear to be of great importance for stabilization of the transporter in its oligomeric state, as we experienced that hNKCC1 only retained its dimeric architecture when solubilized and purified in mild detergents like lauryl maltose neopentyl glycol (LMNG) and glyco-diosgenin (GDN), but not the de-lipidating n-dodecyl-*β*-D-maltopyranoside (DDM). These findings are different from the dimeric KCC transporters that appear to be less lipid-dependent, with DDM routinely used for protein handling without disruption of the oligomeric state (Chi *et al.*, 2021; Liu *et al.*, 2019). This may be a result of the different oligomeric arrangements of the protomers in the transmembrane region of NKCC1 vs KCC structures. The dimers of sodium-independent CCCs rely on stronger interactions between the cytoplasmic domains, with weaker interactions between the transmembrane regions due to a larger distance between the two protomers (Fig. S1C) (Chew *et al.*, 2019; Chi *et al.*, 2021; Xie *et al.*, 2020). Other transporters with low oligomeric stability, like LeuT from *Aquifex aeolicus* or NhaA from *Escherichia coli*, also contain lipids at their dimer interfaces that can easily be broken down by delipidating the protein or mutating residues within the lipid binding sites (Gupta *et al*, 2017). In addition to the crucial interfacial lipids, our cryo-EM map revealed presence of three cholesterol molecules per protomer, all localized at the extracellular part of the cotransporter, locking it in an inward-facing conformation. Interestingly, a similar regulatory role of cholesterol was suggested for the human serotonin transporter and the *Drosophila melanogaster* dopamine transporter (Coleman *et al.*, 2016; Penmatsa *et al.*, 2013), but these two transporters were stabilized in their outward-facing conformations. Cholesterol depletion induced an inward-facing conformation of the serotonin transporter changing the substrate affinity, turnover rate and maximum transport velocity (Laursen *et al*, 2018). Whether cholesterol depletion can provide an experimental approach to generate structural information of NKCC1 in an outward-orientated manner remains to be determined.

Although substrates of the CCC family are much smaller than substrates translocated by other APC transporters, the important coordination networks bridging between the scaffold and bundle domain remain strikingly similar. The Y383 residue on TM3 that is crucial for the LeuT transporter (Zhang *et al.*, 2018) is conserved within the CCC family, with the exception of NCC where it is replaced with a histidine residue that likely undertakes the role of the missing K^+^ substrate. Similar substitutions are observed in, for example, the sodium independent APC transporters where a positively charged basic residue on TM5 replaces the role of the Na^+^ ion or in the case of prokaryotic SLC6 transporters, where a Cl^−^ ion present in eukaryotic transporters is substituted by a glutamate residue on TM7 (Coleman *et al.*, 2016; Penmatsa *et al.*, 2013). The coordination network is important for proper coupling between the main substrate and driving ion and therefore it is often conserved between APC transporters with drastically different substrates. The SLC11 family of divalent metal ions is an example of transporters that do not have this network, resulting in both coupled and uncoupled movements of either of the substrates (Mackenzie *et al*, 2006). A crystal structure of ScaDMT in complex with Mn^2+^ reveals that even though the architecture of the main binding pocket is strikingly similar to the binding pocket of NKCC1, substrate coordination is only maintained by the bundle domain (Ehrnstorfer *et al.*, 2014).

The Na^+^ binding site is highly conserved throughout sodium-dependent APC transporters (Fig. S5A-C) (Faham *et al.*, 2008; Khafizov *et al*, 2012; Krishnamurthy *et al*, 2009; Malinauskaite *et al.*, 2014; Penmatsa *et al.*, 2013; Perez *et al*, 2012; Weyand *et al*, 2008; Yamashita *et al.*, 2005). The crucial role of the Na2 site seems also to be retained in many sodium-independent transporters adopting the LeuT-fold (Fig. S5D-F). Crystal structures revealed that these transporters often contain a charged amine group at this location that undertakes the role of the Na^+^, creating a link between TM1 and TM8 (e.g. R262 of CaiT (Kalayil *et al*, 2013), K158 in ApcT (Shaffer *et al*, 2009) and K154 in BasC (Errasti-Murugarren *et al*, 2019)). Contrastingly, KCC transporters do not contain any basic residues near the Na2 site, thereby suggesting a different mechanism than in other sodium-independent APC transporters. The lack of Na^+^ coupling is reflected in the transport directionality of KCC transporters that translocate a K^+^ and Cl^−^ ion from the cytoplasm to the extracellular space (Fig. 2B and S6A). Structure superposition of hNKCC1 and the hKCC1 revealed a slight 2-3 Å displacement of TM8 that might impact Cl^−^ coupling at the Cl2 site, as KCC transporters only transport one K^+^ and one Cl^−^ (Fig. S7A and B) (Liu *et al.*, 2019). Interestingly, in close proximity to the Na2 site, L297^hNKCC1^ is substituted with Q130^hKCC1^ in KCC1 on TM1, and L616^hNKCC1^ is substituted by Q521^hKCC1^ on TM8. Together with N131^hKCC1^ these residues could create a flexible hydrogen bond network linking TM1 and TM8 in the outward-facing conformation of KCCs that stabilizes the interface between the bundle and scaffold domain (Fig. S6B). These substitutions are not present in any of the sodium-dependent CCC transporters (Fig. S6C).

The Cl2 binding site of hNKCC1 and other CCC’s is placed towards the cytoplasm and bridges between TM10 and the unwound part of TM6, in a similar way as the crucial glutamate residue on TM2 does in SLC6 transporters. For SLC6 type transporters such as LeuT and SERT, the interaction between E62^LeuT^ or the equivalent E136^SERT^ and the unwound junction of TM6 is maintained throughout all the known states, highlighting the importance of this interaction (Focht *et al*, 2021). It seems that the Cl^−^ ion together with Y686^hNKCC1^ replaces the role of E62^LeuT^ and allows for transport translocation. The coordination network between TM2-TM10-Cl^−^-TM6 in NKCC1 therefore appears to play the same role as the TM10-TM2-water-TM6 network within SLC6 transporters connecting the main binding site to distal rearrangements of the transporter. The close interactions between TM10-TM2 and TM6 mediated through a glutamate-water network (SLC6) or a Cl^−^ ion (SLC12) cannot be found in other APC families. Interestingly, structural analysis revealed that other transporters use different interaction types in order to maintain the interactions between these helices. For vSGLT (SLC5) a phenylalanine on TM10 and a second phenylalanine on the unwound part of TM6 make *π*-*π* interactions. Additionally, TM10 and TM2 contain a number of aromatic residues (F447^vSGLT^, Y101^vSGLT^ and F102^vSGLT^) potentially interacting hydrophobically and bringing these two helices in close proximity (Fig. S7 A). In a similar manner the GkApcT (SLC7) transporter contains two aromatic residues maintaining the interaction between TM10 and TM6, whereas the TM2-TM10 interaction is maintained by a salt bridge (Fig. S7 B).

Both Cl^−^ sites of NKCC1 are conserved in KCC transporters, with published KCC structures having both sites occupied (Fig. 5D) (Liu *et al.*, 2019) despite KCCs transporting only one Cl^−^ ion. Interestingly, mutagenesis of residues coordinating either of the two Cl^−^ ions impair transport, suggesting that both Cl^−^ sites are required (Liu *et al.*, 2019). This indicates that the Cl2 site in SLC12 transporters also displays integral, functional properties similar to the conserved glutamate of TM2 in SLC6 transporters (e.g. Glu136^SERT^). We speculate that this site is adapted for a Cl^−^ sensory function in KCC transporters, which activates Cl^−^ efflux at high cytoplasmic concentrations of chloride and oppositely blocks efflux at low cytoplasmic Cl^−^ concentrations.

Structural characterization of hNKCC1 allowed us to generate a homology model of the transmembrane part of hNCC and analyze its ion coupling mechanisms and binding site specificity. As mentioned, NCC contains a histidine residue (H234^NCC^) instead of the crucial tyrosine on TM3 of NKCC and KCC that coordinates the K^+^ ion (Fig. 4C). In its protonated, positively charged state, the histidine residue could replace the K^+^ ion transported by other CCC transporters in a similar manner as a positively charged arginine or lysine residue performs the function of the Na^+^ ion at the Na2 site in sodium-independent APC transporters (Fig. S5D-F). Moreover, the central localization on TM3 would still allow the histidine residue to participate in the important coordination network within the main binding pocket and maintain the high substrate selectivity and coupling. As expected, the Na2 binding site in NCC is also fully conserved compared to NKCC1 and NKCC2 (Fig. 2D), because NCC also uses Na^+^ coupling to drive Cl^−^ movement from the extracellular space to the cytoplasm. Surprisingly, however, sequence alignment and the homology model indicate that both Cl^−^ sites are again conserved (Fig. 5D). It is possible, that two Cl^−^ ions might be required to bind in order to allow the transporter to go through different conformations or Cl^−^ may have an inhibitory effect at high cytoplasmic Cl^−^ concentrations. Interestingly, earlier studies of NKCC1 have provided evidence that the two Cl^−^ binding sites are nonequivalent and exhibit different Cl^−^ affinities: a low Cl^−^ affinity site was identified closer to the cytoplasm and in near proximity of the Na^+^ site (corresponding to the Cl2 site in our structure: K_m_= 55.3 mM) and a high Cl^−^ affinity site found closer to the extracellular environment and in close proximity to the K^+^ site (corresponding to the Cl1 site in our structure: Km=5.1 mM). In addition to the significant difference in Cl^−^ affinity between the two sites, it was also shown that the binding characteristics of the two sites are very different, as Cl2 only was able to bind and transport Cl^−^ ions, whereas Cl1 could also bind and transport NO_3_^−^ and SCN^−^ in addition to Cl^−^ (Brown & Murer, 1985; Kinne *et al*, 1986; Russell, 2000; Turner & George, 1988). We speculate, that if the two predicted Cl^−^ sites in NCC exhibit similar properties as NKCC1, one of the sites might act as a substrate site, whereas the second site with lower Cl^−^ affinity could function as a regulatory site and only allow transport when the cytoplasmic Cl^−^ concentration is low.

For the SLC6 transporters the Na^+^ ion is released first to the intracellular environment followed by the remaining substrates (Li *et al*, 2019; Malinauskaite *et al.*, 2014). Contrastingly, our high-resolution cryo-EM map of hNKCC1 demonstrated solvation of the Cl^−^ ion bound at the Cl2 site, but not of the Na^+^ ion. Independently, our MD simulations of NKCC1 revealed that the Cl^−^ ion is the first to be released, but its release is not a determining step in transport as it is observed to bind reversibly. However, when the Na^+^ ion leaves as a second event in NKCC1 dynamics, none of the released ions rebind, and later the K^+^ and Cl^−^ ion at the main substrate cavity leave and the transporter is ready for return to the outward-open conformation (Fig. 3E). We also identified a possible Na^+^ release pathway along TM5 (Fig. 2E) lined with two glutamate residues at the opening of the cavity that attract and escort the Na^+^ ion to the cytoplasmic environment. Previous reports established that the deactivating phosphorylation of the C-terminus of sodium-independent KCC transporters leads to autoinhibition by blockage of pathway 1 and partially pathway 2 by a 26 residues long part of the flexible N-terminus (Xie *et al.*, 2020). Alternatively, the activating phosphorylation of NKCC1 was hypothesized to interact with the intracellular cytoplasmic domain of the transporter, disrupting a condense, inactive state (Monette & Forbush, 2012; Zhang *et al.*, 2021). We speculate that the lack of TM5 unwinding along pathway 1 together with absence of helix-breaking motifs in the intracellular part of TM5 (Fig. S4), might be replaced with a more pronounced movement of TM1a during substrate release due to activating phosphorylation of the N-terminus.

Interestingly, there is no consensus in the field about the potential substrate release mechanism of CCC transporters. Two different substrate binding and release models were proposed previously (Delpire & Gagnon, 2011; Lytle *et al*, 1998). In the so-called ‘glide-symmetry’ model, the outward-facing conformation of NKCC1 first binds the Na^+^ ion, then the Cl^−^ at the Cl2 site, followed by K^+^ and Cl^−^ from Cl1 site. Then in the inward-facing state, the substrates are released in the same order (Lytle *et al.*, 1998). The ‘steady state’ model, on the other side, suggests that the outward-open state first binds Cl^−^ at the Cl2 site, then the Na^+^ ion, followed by the second Cl^−^ ion at the Cl1 site and the K^+^ ion, whereas the substrates are released a reverse order to the cytoplasmic environment (Delpire & Gagnon, 2011). The substrate release mechanism proposed in our work does not fit any of the two hypothesized substrate release mechanisms. Intracellular release of Cl^−^ from Cl2 prior to the Na^+^ ion was independently observed, first by visualization of the intracellular solvation network of hNKCC1 leading to hydration of the Cl^−^ but not the Na^+^ ion, and secondly by use of MD simulations where the Cl^−^ ion was shown to leave the transporter before the Na^+^ ion. We speculate that the observed reversibility of Cl^−^ binding and unbinding at Cl2 found in our MD simulations might have prevented proper evaluation of the substrate release order in the previously published studies. Interestingly, if the ‘steady state’ model assumed glide-symmetry instead of mirror-symmetry, as found for other LeuT-fold transporters, the mechanism would fit well with our findings. However, we cannot exclude that our release model might have been impacted by limitations of the cryo-EM approach applied. For example, extraction of the transporter from the native membrane followed by detergent-based protein purification changes the system significantly when compared to the natural environment of NKCC1 in cells. Furthermore, a lack of other cell components, protein interaction partners, alterations in fluid composition (needed to stabilize the isolated protein) or a lack of membrane potential might have a pronounced effect on the transport dynamics. Therefore, further structures of the CCCs in different conformational states are still needed to properly understand the substrate binding and release events.

In conclusion, SLC12 transporters are of great importance for proper functioning of all organisms ranging from humans and vertebrate animals through plants (Colmenero-Flores *et al*, 2007) and fungi to insects, worms and cyanobacteria (Hartmann *et al*, 2014). Insight into the three-dimensional structure and mechanistic details of substrate binding, translocation and release might lead to better understanding of the role of these transporters in normal physiology and their pathophysiological roles. Our 2.6Å structure of occluded and inward-facing hNKCC1 provides not only a direct insight into substrate accommodation, but it is the first structure of a SLC12 transporter with a high enough resolution to resolve the intracellular water networks and reveal solvation of Cl^−^ at Cl2 but not Na^+^ (Fig. 2A and 4A). By a combination of structural studies and MD simulations we show that the Cl^−^ ion is released before the Na^+^ ion and we also identify a potential intracellular release pathway of Na^+^ lined by two glutamate residues crucial for attracting and escorting the cation out of the transporter. Finally, we identify the Cl2 site as a potential sensory and stabilizing site of the transporter, where the Cl^−^ has the same role as a strictly conserved glutamate residue in SLC6 transporters, maintaining the coupling network between the unwound part of TM6 and distal parts of the transporter to enable conformational transitions of NKCC1. In general, insight into the ion coupling networks and the transport mechanism of the CCCs can improve our understanding of not only the physiological roles of the transporter and other SLC12 transporters, but also help with determination of specific buffer composition, mutations and/or nanobodies necessary for structural studies focusing on different conformational states of the CCCs, e.g. in a similar manner as earlier done for LeuT (Focht *et al.*, 2021; Gotfryd *et al.*, 2020; Krishnamurthy & Gouaux, 2012; Malinauskaite *et al.*, 2016). In addition, knowledge of the exact molecular contributions of significant residues during the transport cycle can further help with rational drug design, thereby leading to safer and more target-specific pharmaceuticals, in contrary to the less-specific loop diuretics used nowadays to treat hypertension and edema (e.g. bumetanide inhibits NKCC1, NKCC2 as well as KCC1-4) (Gillen *et al*, 1996; Holtzman *et al*, 1998; Race *et al*, 1999).

## Materials and methods

### Reagents

All chemicals were obtained from Sigma, unless stated otherwise.

### Expression and purification of the human NKCC1

The full-length human NKCC1 with a N-terminal 8xHis-TwinStrep-GFP-tag-3Cprotease_cleavage_site was expressed in HEK293 GnTl^-^ (ATCC) cells by use of the BacMam system as described earlier (Goehring *et al*, 2014). HEK293 GnTl^-^ cells were grown to a density of 3 × 10^6^ cells/mL at 37°C and 5% CO_2_ on an orbital shaker at 120 rpm. The cells were then transduced with the P3 NKCC1 BacMam virus to a MOI of 3 and the cells were further incubated in the incubator. 24h post transduction 10mM sodium butyrate was added to the cells and the temperature was lowered to 30°C with 5% CO_2_ while shaking (120 rpm). After 48h the cells were harvested by centrifugation at 6,200 × g at 4°C. The cell pellets were stored at −80°C.

The frozen cell pellet was thawed and resuspended in a buffer composed of 200mM NaCl, 200mM KCl, 20mM Tris-HCl pH 8.0, 10% (v/v) glycerol and 0.07mM bumetanide. The cells lysed by sonication and centrifugated for 20 min at 20,000 × g at 4°C. The collected supernatant was further centrifugated at 163,000 × g for 3 hours at 4°C and the pellet was thereafter resuspended in the same buffer with a ratio of 10mL buffer per 1g of wet pellet. 1% lauryl maltose neopentyl glycol (LMNG, Anatrace) and 0.1% cholesteryl hemisuccinate (CHS, Anatrace) was added and the membrane proteins were solubilized overnight at 4°C. The mixture was then centrifugated at 14000 × g for 30 min, the insoluble fraction was discarded and the supernatant was incubated with Strep-Tactin resin for 3h. The resin was washed with 10 column volumes washing buffer containing 0.01% LMNG, 0.001% CHS, 200mM NaCl, 200mM KCl, 20mM Tris-HCl pH 8.0, 10% (v/v) glycerol and 0.07mM bumetanide. The hNKCC1 protein was eluted with wash buffer supplemented with 5mM desthiobiotin. The sample was then cleaved overnight with 3C protease and concentrated to 3 mg/mL. The protein was further purified by gel filtration (Superose 6 3.2/300) on an Äkta Purifier system equilibrated in a buffer composed of 0.01% glycol-diosgenin (GDN, Anatrace), 200mM NaCl, 200mM KCl, 20mM Tris-HCl pH 8.0 and 0.07mM bumetanide. Peak fractions were collected and concentrated to 0.9-1.5mg/mL for cryo-EM studies.

### Thallium ion (Tl^+^) flux transport assay

The NKCC1 mutant constructs were designed in the same way as the wt NKCC1 construct used for structure determination. The mutants were prepared by GenScript. Activity levels of the wild-type NKCC1 transporter and NKCC1 mutants were determined by use of the thallium ion flux transport assay. The construct with a N-terminal 8xHis-TwinStrep-GFP-tag were expressed in GripTite HEK293 MSR cells (Thermo Fisher) in black 96 well plates with transparent bottoms. The cells were maintained in DMEM GlutaMAX medium (Gibco) with addition of 10% FBS and 1x non-essential amino acid (NEAA) cell culture supplement. 50,000 cells were seeded out per well and the cells were incubated for 24h at 37° with 8% CO_2_. On the day of transfection, 3 μL Lipofectamine 3000 (Invitrogen) was mixed with 47 μL Opti-Mem medium (Gibco). Separately, 1000ng DNA was diluted in 36 μL Opti-MEM medium with addition of 4 μL P3000 reagent (Invitrogen) and thereafter, the Lipofectamine 3000 solution was mixed with the DNA solution. After 15 min. incubation, 10 μL of the mixture was added to each of the wells. After 5-6 hours, 100 μL DMEM medium, 1x NEAA and 30% FBS was added to each well. The activity assay was performed 72 h post transfection of the cells.

On the day of the experiment, the DMEM medium was exchanged for 80 μL loading buffer containing probenecid, PowerLoad and FluxOR II Reagent from the FluxOR II Red Potassium Ion Channel Assay kit (Thermo Fisher Scientific) and a low chloride buffer (15 mM Na-HEPES pH 7.4, 135 mM Na-gluconate, 1 mM MgCl_2_-6H_2_O, 1 mM Na_2_SO_4_ and 1 mM CaCl_2_-2H_2_O). The cells were incubated with the loading buffer for 1 h at room temperature and thereafter, they were washed once with the low chloride buffer with addition of probenecid, followed by addition of 80 μL buffer to each well.

The assay was performed using the ENSPIRE 2300 kinetic dispense microplate reader (PerkinElmer). First, GFP fluorescence was measured for later normalization (excitation at 480 nm, emission at 510 nm, 100 flashes, 3 repeats, 0.1s). After 5 s of baseline recording (10 repeats) the transport assay was initiated by addition of 20 μL stimulation buffer containing low chloride buffer, 135 mM NaCl, and 5 mM thallium sulfate with or without 165 μM bumetanide (final concentrations of 27 mM NaCl, 1 mM thallium sulfate and 33 μM of bumetanide). The excitation wavelength was set to 565nm and the emission wavelength was set to 583nm. The plate was read every 0.5s for 80 repeats (40s).

### Biotinylation assay

Cell surface protein biotinylation was performed to estimate the plasma membrane expression of various NKCC1 mutants relative to the wild-type transporter. The N-terminal 8xHis-TwinStrep-GFP-tagged NKCC1 constructs were expressed in GripTite HEK293 MSR cells (Thermo Fisher) in 24 well plates. The transfection of cells was performed in the same was as done for the Tl^+^ uptake assay, but the amounts were scaled up. The biotinylation assay was performed 72 h after transfection of the cells.

The cells were washed in basic buffer (135 mM NaCl, 5 mM KCl, 1 mM CaCl2-2H20, 1 mM MgCl2-6H20, 1 mM Na2HPO4-2H20, 1 mM Na2SO4, 15 mM Sodium HEPES pH 7.4) and incubated in the low chloride buffer (same as in uptake assay) for 1 h at 18-24°C. Thereafter, the cells were placed on ice and washed with ice-cold PBS-CM buffer (PBS pH 7.5 with addition of 1 mM CaCl_2_, 0.1 mM MgCl_2_), followed by the biotinylation buffer (10 mM triethanolamine, 2 mM CaCl_2_, 125 mM NaCl). The cells were incubated in the biotinylation buffer with addition of 1 mg/mL of sulfosuccinimidyl 2-(biotin-amido)-ethyl-1,3-dithiopropionate (EZ-link Sulfo-NHS-SS-biotin, Pierce) for 30 min at 4°C. After the incubation period, the cells were washed with ice-cold quenching buffer (50 mM Tris-HCl in PBS-CM buffer), followed by the PBS-CM buffer. The cells were incubated in lysis buffer (20 mM Tris-HCl pH 8.0, 5 mM EDTA, 150 mM NaCl, 1% Triton X-100, 0.2% BSA and Halt® protease inhibitors) for 30 min on ice, followed by probe sonication. Thereafter, the homogenates were centrifugated at 10,000xg for 5 minutes at 4°C. One fraction was retained for total NKCC1 protein abundance estimation and the remainder was incubated for 1 h with Neutravadin resin at room temperature under rotation. The resin was washed four times with PBS containing protease inhibitors and the bound protein was eluted by addition of SDS-PAGE sample buffer to the resin. Western blotting was performed using standard methods using anti-NKCC1 antibody.

### Cryo-EM sample preparation and data collection

For preparation of grids, freshly purified hNKCC1 protein was used. Grids used were UltrAuFoil 1.2/1.3-300, glow-discharged in residual atmospheric air for 45 sec at 15 mA in a GloQube (Quorum). A 3 uL drop was applied to the gold foil side of the grid and blotted in a Vitrobot Mark IV (ThermoFisher Scientific) using a blot force of 0 and blot time of 3-4 seconds before plunge-freezing into liquid ethane cooled by liquid nitrogen. Micrograph data was collected on a Titan Krios G3i microscope (ThermoFisher Scientific) operated at 300 KeV equipped with a BioQuantum energy filter (energy slit width 20 eV) and K3 camera (Gatan). For data sets 1 and 2 (collected at the Krios2 microscope, EMBION facility, Aarhus University) a nominal magnification of 165,000x was used, resulting in a physical pixel size of 0.507 Å^2^/px with a total dose of 40.12 e^−^/Å^2^ for data set 1 and a total dose of 39.78 e^−^/Å^2^ for data set 2. For both data sets, movies were fractionated into 43 frames. The defocus range was set to 0.6-1.6 micron. For dataset 3 (collected on Titan Krios at eBIC, Oxford, UK), a nominal magnification of 130,000x was used, resulting in a physical pixel size of 0.83 Å^2^/px, with movies saved in super-resolution pixel size of 0.415 Å^2^/px. A total dose of 60.438 e^−^/Å^2^ per movie was spread across 40 frames. The defocus range was set to 0.5-2.4 micron.

### Cryo-EM data processing

Datasets 1 and 2 were processed exclusively in RELION-3.1 (Scheres, 2012) (Fig. EV1), whereas dataset 3 was processed primarily in RELION-3.1 with some classifications performed using the heterogeneous refinement in cryosparc3 (Punjani *et al.*, 2017) (Fig. EV2). For all data sets, movies were motion corrected using RELION’s own MotionCorr implementation and CTFFind4 (Rohou & Grigorieff, 2015) used for determination of CTF parameters. Only micrographs with CTF fit resolution better than 4 Å were used onwards. Initial particle picking was done using RELION’s LoG-picker on a small, random subset of micrographs in data set 1. Picked particles were extracted with a 512-pixel box, downsampled to 128 pixels, and subjected to 2D classification. Clear junk particles were discarded, and the remainder of particles were used in 3D auto-refinement with the zNKCC1 reconstruction (EMD-0473) as initial reference, lowpass filtered to 30 Å resolution. The resulting 3D reconstruction was used in 2D classification without image alignment, and the best 2D class averages were used for template-based particle picking on all micrographs for all three data sets. Particle images were extracted in a 480-pixel box, downsampled to 128 pixels, and 2D classified using fast subsets and skipping CTF until first peak. Clear junk classes were discarded, and the remainder raw particle set was subjected to 3D classification using fast subsets and 8 classes for data set 1 and 10 classes for data set 2. Particles from the 3D classes displaying features of an intact detergent micelle were selected and run through 3D auto-refinement followed by 2D classification without image alignment. Again, clear junk was discarded, and the remainder particles were re-extracted with downsampling to a 240-pixel box (1.014 Å^2^/px). Particles were again subjected to 3D classification now without fast subsets, using 8 and 6 classes for data set 1 and 2, respectively. The 3D classes displaying clear density features of transmembrane helices were selected and subjected to unmasked 3D auto-refinement. The resulting consensus reconstructions were deemed of good enough quality to proceed with Bayesian polishing, followed by 3D auto-refinement with a mask around the transmembrane domains and application of C2 symmetry. The resulting masked 3D reconstructions were resolved better than 3 Å, which upon merging of the particles from data sets 1 and 2 were 3D auto-refined to a high-quality reconstruction of 2.6 Å resolution. For data set 3, subsequent to 2D classification and the first 3D classification in RELION-3.1, the particle image stack was further cleaned by discarding junk classes from a 2D classification job without image alignment. The resulting stack of 595,171 particle images were re-extracted in a 384-pixel box without downsampling before loaded into Cryosparc3 and further classified by ab-initio reconstruction and heterogenous refinement using three classes. A single class of particles showed high-resolution features for the transmembrane helices, with the cytoplasmic domains being almost impossible to see. This new stack of particle images was taken back into RELION-3.1, cleaned yet again using 2D classification without image alignment, and particle image re-extracted in a 294-pixel box with downsampling to 240 pixels (1.016 Å^2^/px). The now clean stack of particle images was subjected to 3D auto-refinement, Bayesian polishing and 3D auto-refinement again using a mask around the transmembrane domains and applying C2 symmetry. The resulting reconstruction showed high-quality density features resolved slightly better than 3 Å. Using CTF refinement for anisotropic magnification to account for the small discrepancy in pixel size, the data set 3 particles were merged with the already combined particle data from data set 1 and 2, to give a final 3D-refined reconstruction of 2.55-Å average resolution for the transmembrane domains (Fig. EV2A). Local resolution estimation was done using RELION’s own implementation (Fig. EV2B).

### Model building and refinement

The transmembrane domain of hNKCC1 was built in Coot (Emsley *et al*, 2010), guided by the *Danio rerio* NKCC1 structure (PDB entry: 6NPL) (Chew *et al.*, 2019). The refinement was done by use of Real Space Refinement in Phenix (Afonine *et al*, 2018) and manual corrections of the structure were performed in Coot. Validation of the structure was done in MolProbity (Williams *et al*, 2018).

### System setup for molecular dynamics simulations

A model of hNKCC1 was constructed using the Cryo-EM structure of the transmembrane domain of hNKCC1 in complex with its substrates, one Na^+^, one K^+^ and two Cl^−^ ions. In addition, four CHS molecules identified in the Cryo-EM structure were replaced by the cholesterol molecules. The model was subsequently embedded into a flat, mixed lipid bilayer consisting of POPC/POPE/cholesterol at a 2:2:1 ratio, and solvated in a cubic water box containing 0.1M NaCl and 0.1M KCl. The size of the box was 13.1 nm, 13.1 nm and 10.2 nm in the x, y and z dimension, respectively, resulting in ~180,000 atoms in total. The CHARMM36m force field was used for the protein and the CHARMM36 lipid force field was used for all lipid molecules.

In the MD simulations, the temperature was kept constant at 310 K using a Nose-Hoover thermostate with a 1 ps coupling constant, and the pressure at 1.0 bar using the Parrinello-Rahman barostat with a 5 ps time coupling constant. A cutoff of 1.2 nm was applied for the van der Waals interactions using a switch function starting at 1.0 nm. The cutoff for the short-range electrostatic interactions was also at 1.2 nm and the long-range electrostatic interactions were calculated by means of the particle mesh Ewald decomposition algorithm with a 0.12 nm mesh spacing. A reciprocal grid of 112 x 112 x 96 cells was used with 4th order B-spline interpolation. All MD simulations were performed using Gromacs2019.6 (Abraham *et al*, 2015). Twenty independent simulations (200 ns for each) were performed for the hNKCC1 model, so resulting in 8 μs simulations (20 runs × 200 ns × 2 monomers) in total.

## Acknowledgements

The authors are grateful to technical assistance by Tina Drejer, Tetyana Klymchuk, Anna Marie Nielsen, Bente Andersen and Anne Lillevang. We want to thank Thomas Boesen, Andreas Bøggild and Taner Drace from the cryo-EM facility at Aarhus University for assistance with grid screening and numerous overnight data collections. We are thankful to Milena Timcenko, Jeppe Achton Nielsen, Søren Kirk Amstrup and Jonathan Juhl for help with data processing and fruitful discussions about cryo-EM. We want to acknowledge Qi Wu for performing MS on the purified hNKCC1 sample in order to assure that sample of the expressed and purified hNKCC1 construct was correct.

We acknowledge Diamond Light Source for access and support of the cryo-EM facilities at the UK’s National Electron Bio-Imaging Centre (eBIC) under proposal AP27 funded by the Wellcome Trust, MRC and BBRSC.

Work on the project was supported by a PhD fellowship from the Lundbeck Foundation to CN (2015-3225), the Leducq Foundation (17CVD05), the Novo Nordisk Foundation (NNF21OC0067647, NNF17OC0029724, NNF19OC0058439) and the Independent Research Fund Denmark to RF and from the Brainstruc centre (R155-2015-2666 and R328-2019-546) and a professorship grant (R310-2018-3713) funded by the Lundbeck Foundation to PN.

## Author contributions

PN and RF conceived the project, JAL designed the expression constructs and transient transfection screens with CN, and CN established the BacMam expression system with support and guidance of HHG and RH. CN expressed and purified the protein with help of MH, JAL and PN. RKF also purified the protein. CN, RKF and JAL prepared cryo-EM grids, CN and RKF performed data processing, model refinement and validation with support and advice of JLK and JAL. The uptake assay was set-up and optimized by LLR and the final thallium uptake experiments of the wt NKCC1 and mutants were performed by CN under guidance of LLR and RF. Molecular dynamics simulations were done by YW and KLL. The manuscript was drafted by CN. All authors commented on the manuscript.

## Extended View Figures

**Fig. EV1 Processing of cryo-EM datasets 1 and 2 in Relion.** Representative cryo-EM micrographs and two-dimensional class averages are shown for both datasets as well as the processing strategies used to obtain a 2.6Å structure of the transmembrane domain of hNKCC1.

**Fig. EV2 Processing of the third cryo-EM dataset of hNKCC1 in Relion and CryoSPARC3.** Representative cryo-EM micrographs, two-dimensional class averages and the processing strategy used to obtain the final structure of the transmembrane domain of hNKCC1.

**Fig. EV3 Cryo-EM structure of hNKCC1 in complex with sodium, potassium and chloride ions.** A) Gold-standard Fourier shell correlation curves between the two half maps with the final resolution at F=0.143. B) Local resolution estimations throughout the determined transmembrane domain of hNKCC1. C) Cryo-EM map of the cytoplasmic domain of hNKCC1 with the *Danio rerio* NKCC1 model docked (PDB: 6NPJ). D) Representative densities of the different transmembrane helices of molecule A of hNKCC1.

**Fig. EV4 Expression, purification and functional studies of hNKCC1.** A) Purification of hNKCC1 transporter expressed in HEK293 GnTl^−^ cells. The protein used for structure determination was purified by Strep-Tactin purification, Ni^2+^ - ion affinity chromatography (SDS-PAGE 1) followed by size exclusion chromatography (SDS-PAGE 2 and size exclusion chromatogram). B) Initial rate of Tl^+^ transport in non-transfected cells (with and without bumetanide) as well as in cells transfected with wild-type hNKCC1 (with and without bumetanide). C) Representative western blotting of NKCC1 in total and biotinylated (plasma membrane) fractions of wt NKCC1 and mutant NKCC1 expressing cells. D) Surface expression of NKCC1 mutants relative to wt NKCC1.

## Supplementary Figures

**Fig. S1 Comparison of the hNKCC1 structure with previously published SLC12 structures.** A) Comparison of protomer A of the determined hNKCC1 structure (purple) with other published NKCC1 structures (PDB codes: 6NPH, 6PZT; presented in grey) or B) with published KCC structures (PBD codes: 6KKR, 6M1Y, 6UKN, 7D14; presented in grey); C) Comparison of the determined hNKCC1dimer (cyan) with the KCC dimer (grey). D), E), F) Density for potential cholesterol binding site 1, 2, 3 and 4 in the published KCC3 structure (PDB entry: 6M22).

**Fig. S2 Substrate binding of SLC12 and SLC6 transporters.** Similar substrate binding in the interface between the scaffold (wheat) and bundle domain (cyan) of A) NKCC1 belonging to the SLC12 family and B) MhsT belonging to the SLC6 family.

**Fig. S3 Comparison of the cryo-EM structure of hNKCC1 in the occluded, inward-facing conformation with LeuT and MhsT structures in different conformational states.** A) Structural comparison of helices comprising the intracellular vestibule of the occluded, inward-facing conformation of hNKCC1 with the occluded, outward-facing conformation of LeuT (PDB entry: 2A65), the occluded, inward-facing state of MhsT with an unwound TM5 (PDB entry: 4US3; the structure was crystallized by use of the HiLiDe method), the occluded, inward-facing state of MhsT with a refolded TM5 (PDB entry: 4US4; the structure was crystallized by use of the LCP method) and the apo inward open structure of LeuT (PDB entry: 3TT3). Interactions between TM1a and TM5 are visualized. The hNKCC1 structure is shown in gold, whereas the other structures are presented in grey. B) The Na2 sodium binding site of the occluded, outward-facing conformation of LeuT, the occluded, inward-facing state of MhsT with an unwound TM5, the occluded, inward-facing state of MhsT with a refolded TM5 and the apo inward open structure of LeuT.

**Fig. S4 Absence of helix-breaking motifs in CCC transporters.** Sequence alignment of the intracellular part of TM5 containing the GlyX_9_Pro motif in the SLC6 family with sequences of TM5 in CCC transporters.

**Fig. S5 Na2 binding site in sodium-dependent and sodium (A-C) and sodium independent (D-E) APC transporters.** A) NSS family: MhsT (occluded inward-facing, PDB: 4US3), B) SSS family: vSGLT (inward-open, PDB: 3DH4) C) NCS1 family: Mhp1 (occluded ligand bound, PDB: 4D1A), D) BCCT family: CaiT (inward-facing, 2WsW), E) APC family: ApcT (inward-open, 3GI9), F) LAT family: BasC (inward-open, 6F2W).

**Fig. S6 Lack of a sodium binding site in KCC transporters.** A) Superposition of TM1 and TM8 in NKCC1 and KCC1, B) Comparison of Na2 binding site and the part below the Na2 binding site of NKCC1 and KCC1, C) Sequence alignment of the Na2 binding site and the fragment below the Na2 binding site of the CCC transporters. The G-A substitution of KCC1-4 on TM8 responsible for abolishment of sodium binding is highlighted as well as the two Q residues on TM1 and TM8 that are a part of the hydrogen binding network holding TM1-TM8 together.

**Fig. S7 Coordination network between TM2-TM10 and the unwound part of TM6 in transporters belonging to other SLC families** A) vSGLT (SLC5, 3DH4) and B) GkApcT (SLC7, 5OQT).

## Tables

**Table 1.**
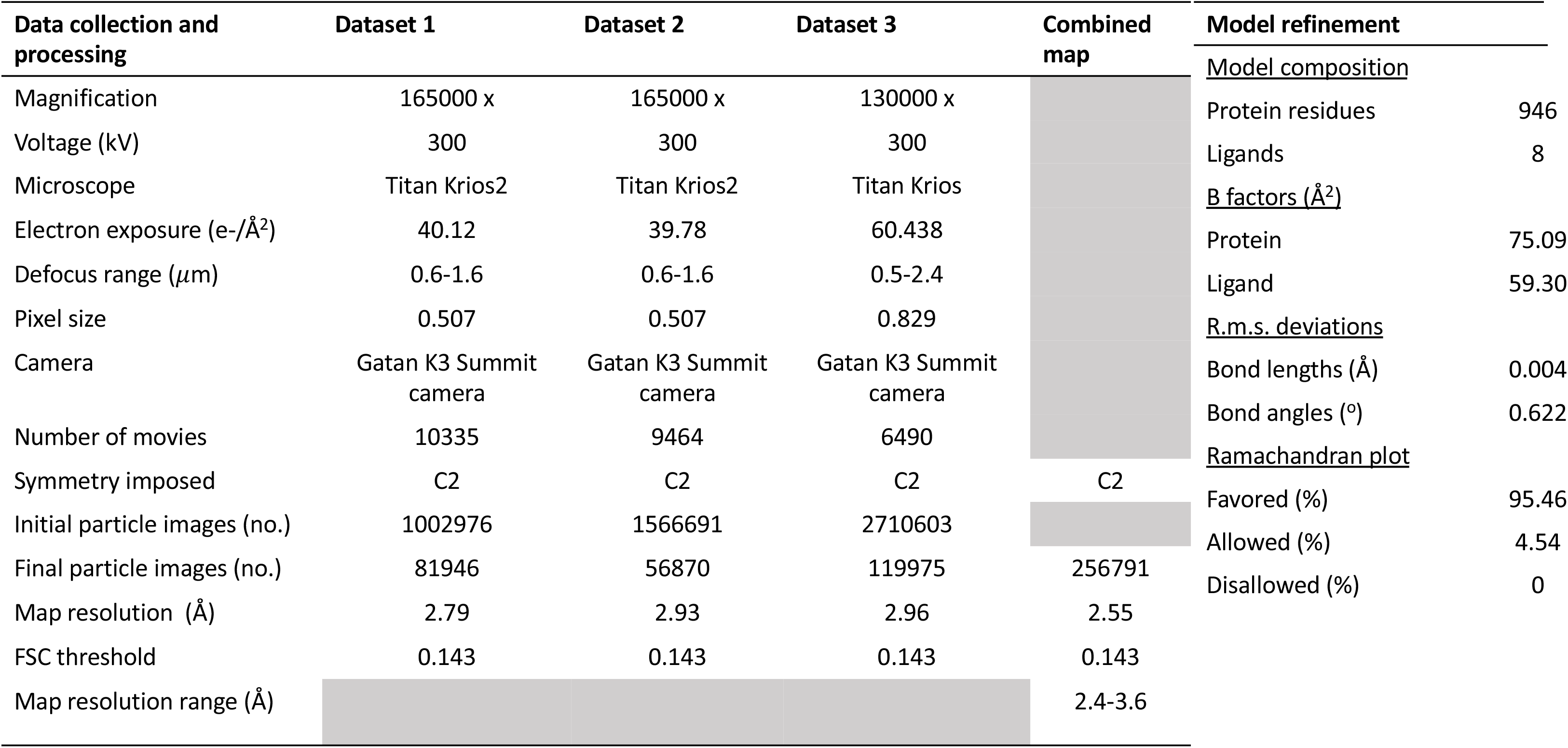
Cryo-EM data collection, refinement and validation.

## References

Abraham JA, Murtola T, Schulz R, Pall S, Smith JC, Hess B, Lindahl E (2015) GROMACS: High performance molecular simulations through multi-level parallelism from laptops to supercomputers. SoftwareX 1-2: 19–25

Afonine PV, Poon BK, Read RJ, Sobolev OV, Terwilliger TC, Urzhumtsev A, Adams PD (2018) Real-space refinement in PHENIX for cryo-EM and crystallography. Acta Crystallogr D Struct Biol 74: 531–544

Arroyo JP, Kahle KT, Gamba G (2013) The SLC12 family of electroneutral cation-coupled chloride cotransporters. Mol Aspects Med 34: 288–298

Ben-Yona A, Kanner BI (2009) Transmembrane domain 8 of the {gamma}-aminobutyric acid transporter GAT-1 lines a cytoplasmic accessibility pathway into its binding pocket. J Biol Chem 284: 9727–9732

Berka K, Hanak O, Sehnal D, Banas P, Navratilova V, Jaiswal D, Ionescu CM, Svobodova Varekova R, Koca J, Otyepka M (2012) MOLEonline 2.0: interactive web-based analysis of biomacromolecular channels. Nucleic Acids Res 40: W222–227

Blaesse P, Airaksinen MS, Rivera C, Kaila K (2009) Cation-chloride cotransporters and neuronal function. Neuron 61: 820–838

Brown CD, Murer H (1985) Characterization of a Na: K: 2C1 cotransport system in the apical membrane of a renal epithelial cell line (LLC-PK1). J Membr Biol 87: 131–139

Chen N, Vaughan RA, Reith ME (2001) The role of conserved tryptophan and acidic residues in the human dopamine transporter as characterized by site-directed mutagenesis. J Neurochem 77: 1116–1127

Chew TA, Orlando BJ, Zhang J, Latorraca NR, Wang A, Hollingsworth SA, Chen DH, Dror RO, Liao M, Feng L (2019) Structure and mechanism of the cation-chloride cotransporter NKCC1. Nature 572: 488–492

Chi X, Li X, Chen Y, Zhang Y, Su Q, Zhou Q (2020) Cryo-EM structures of the full-length human KCC2 and KCC3 cation-chloride cotransporters. Cell Res

Chi X, Li X, Chen Y, Zhang Y, Su Q, Zhou Q (2021) Cryo-EM structures of the full-length human KCC2 and KCC3 cation-chloride cotransporters. Cell Res 31: 482–484

Claxton DP, Quick M, Shi L, de Carvalho FD, Weinstein H, Javitch JA, McHaourab HS (2010) Ion/substrate-dependent conformational dynamics of a bacterial homolog of neurotransmitter:sodium symporters. Nat Struct Mol Biol 17: 822–829

Coleman JA, Green EM, Gouaux E (2016) X-ray structures and mechanism of the human serotonin transporter. Nature 532: 334–339

Colmenero-Flores JM, Martinez G, Gamba G, Vazquez N, Iglesias DJ, Brumos J, Talon M (2007) Identification and functional characterization of cation-chloride cotransporters in plants. Plant J 50: 278–292

Delpire E, Gagnon KB (2011) Kinetics of hyperosmotically stimulated Na-K-2Cl cotransporter in Xenopus laevis oocytes. Am J Physiol Cell Physiol 301: C1074–1085

Delpire E, Gagnon KB (2018) Na(+) -K(+) -2Cl(-) Cotransporter (NKCC) Physiological Function in Nonpolarized Cells and Transporting Epithelia. Compr Physiol 8: 871–901

Demian WL, Persaud A, Jiang C, Coyaud E, Liu S, Kapus A, Kafri R, Raught B, Rotin D (2019) The Ion Transporter NKCC1 Links Cell Volume to Cell Mass Regulation by Suppressing mTORC1. Cell Rep 27: 1886–1896 e1886

Doyle DA, Morais Cabral J, Pfuetzner RA, Kuo A, Gulbis JM, Cohen SL, Chait BT, MacKinnon R (1998) The structure of the potassium channel: molecular basis of K+ conduction and selectivity. Science 280: 69–77

Ehrnstorfer IA, Geertsma ER, Pardon E, Steyaert J, Dutzler R (2014) Crystal structure of a SLC11 (NRAMP) transporter reveals the basis for transition-metal ion transport. Nat Struct Mol Biol 21: 990–996

Emsley P, Lohkamp B, Scott WG, Cowtan K (2010) Features and development of Coot. Acta Crystallogr D Biol Crystallogr 66: 486–501

Errasti-Murugarren E, Fort J, Bartoccioni P, Diaz L, Pardon E, Carpena X, Espino-Guarch M, Zorzano A, Ziegler C, Steyaert J et al (2019) L amino acid transporter structure and molecular bases for the asymmetry of substrate interaction. Nat Commun 10: 1807

Faham S, Watanabe A, Besserer GM, Cascio D, Specht A, Hirayama BA, Wright EM, Abramson J (2008) The crystal structure of a sodium galactose transporter reveals mechanistic insights into Na+/sugar symport. Science 321: 810–814

Fenollar-Ferrer C, Stockner T, Schwarz TC, Pal A, Gotovina J, Hofmaier T, Jayaraman K, Adhikary S, Kudlacek O, Mehdipour AR et al (2014) Structure and regulatory interactions of the cytoplasmic terminal domains of serotonin transporter. Biochemistry 53: 5444–5460

Focht D, Neumann C, Lyons J, Eguskiza Bilbao A, Blunck R, Malinauskaite L, Schwarz IO, Javitch JA, Quick M, Nissen P (2021) A non-helical region in transmembrane helix 6 of hydrophobic amino acid transporter MhsT mediates substrate recognition. EMBO J 40: e105164

Forrest LR, Kramer R, Ziegler C (2011) The structural basis of secondary active transport mechanisms. Biochim Biophys Acta 1807: 167–188

Forrest LR, Rudnick G (2009) The rocking bundle: a mechanism for ion-coupled solute flux by symmetrical transporters. Physiology (Bethesda) 24: 377–386

Gillen CM, Brill S, Payne JA, Forbush B, 3rd (1996) Molecular cloning and functional expression of the K-Cl cotransporter from rabbit, rat, and human. A new member of the cation-chloride cotransporter family. J Biol Chem 271: 16237–16244

Goehring A, Lee CH, Wang KH, Michel JC, Claxton DP, Baconguis I, Althoff T, Fischer S, Garcia KC, Gouaux E (2014) Screening and large-scale expression of membrane proteins in mammalian cells for structural studies. Nat Protoc 9: 2574–2585

Gotfryd K, Boesen T, Mortensen JS, Khelashvili G, Quick M, Terry DS, Missel JW, LeVine MV, Gourdon P, Blanchard SC et al (2020) X-ray structure of LeuT in an inward-facing occluded conformation reveals mechanism of substrate release. Nat Commun 11: 1005

Gupta K, Donlan JAC, Hopper JTS, Uzdavinys P, Landreh M, Struwe WB, Drew D, Baldwin AJ, Stansfeld PJ, Robinson CV (2017) The role of interfacial lipids in stabilizing membrane protein oligomers. Nature 541: 421–424

Hartmann AM, Blaesse P, Kranz T, Wenz M, Schindler J, Kaila K, Friauf E, Nothwang HG (2009) Opposite effect of membrane raft perturbation on transport activity of KCC2 and NKCC1. J Neurochem 111: 321–331

Hartmann AM, Nothwang HG (2014) Molecular and evolutionary insights into the structural organization of cation chloride cotransporters. Front Cell Neurosci 8: 470

Hartmann AM, Tesch D, Nothwang HG, Bininda-Emonds OR (2014) Evolution of the cation chloride cotransporter family: ancient origins, gene losses, and subfunctionalization through duplication. Mol Biol Evol 31: 434–447

Hebert SC, Mount DB, Gamba G (2004) Molecular physiology of cation-coupled Cl- cotransport: the SLC12 family. Pflugers Arch 447: 580–593

Holtzman EJ, Kumar S, Faaland CA, Warner F, Logue PJ, Erickson SJ, Ricken G, Waldman J, Kumar S, Dunham PB (1998) Cloning, characterization, and gene organization of K-Cl cotransporter from pig and human kidney and C. elegans. Am J Physiol 275: F550–564

Jaggi AS, Kaur A, Bali A, Singh N (2015) Expanding Spectrum of Sodium Potassium Chloride Co-transporters in the Pathophysiology of Diseases. Curr Neuropharmacol 13: 369–388

Ji W, Foo JN, O’Roak BJ, Zhao H, Larson MG, Simon DB, Newton-Cheh C, State MW, Levy D, Lifton RP (2008) Rare independent mutations in renal salt handling genes contribute to blood pressure variation. Nat Genet 40: 592–599

Joseph D, Pidathala S, Mallela AK, Penmatsa A (2019) Structure and Gating Dynamics of Na(+)/Cl(-) Coupled Neurotransmitter Transporters. Front Mol Biosci 6: 80

Jungnickel KEJ, Parker JL, Newstead S (2018) Structural basis for amino acid transport by the CAT family of SLC7 transporters. Nat Commun 9: 550

Kalayil S, Schulze S, Kuhlbrandt W (2013) Arginine oscillation explains Na+ independence in the substrate/product antiporter CaiT. Proc Natl Acad Sci U S A 110: 17296–17301

Kazmier K, Sharma S, Quick M, Islam SM, Roux B, Weinstein H, Javitch JA, McHaourab HS (2014) Conformational dynamics of ligand-dependent alternating access in LeuT. Nat Struct Mol Biol 21: 472–479

Khafizov K, Perez C, Koshy C, Quick M, Fendler K, Ziegler C, Forrest LR (2012) Investigation of the sodium-binding sites in the sodium-coupled betaine transporter BetP. Proc Natl Acad Sci U S A 109: E3035–3044

Kinne R, Kinne-Saffran E, Scholermann B, Schutz H (1986) The anion specificity of the sodium-potassium-chloride cotransporter in rabbit kidney outer medulla: studies on medullary plasma membranes. Pflugers Arch 407 Suppl 2: S168–173

Korkhov VM, Holy M, Freissmuth M, Sitte HH (2006) The conserved glutamate (Glu136) in transmembrane domain 2 of the serotonin transporter is required for the conformational switch in the transport cycle. J Biol Chem 281: 13439–13448

Koumangoye R, Bastarache L, Delpire E (2021) NKCC1: Newly Found as a Human Disease-Causing Ion Transporter. Function (Oxf) 2: zqaa028

Krishnamurthy H, Gouaux E (2012) X-ray structures of LeuT in substrate-free outward-open and apo inward-open states. Nature 481: 469–474

Krishnamurthy H, Piscitelli CL, Gouaux E (2009) Unlocking the molecular secrets of sodium-coupled transporters. Nature 459: 347–355

Laursen L, Severinsen K, Kristensen KB, Periole X, Overby M, Muller HK, Schiott B, Sinning S (2018) Cholesterol binding to a conserved site modulates the conformation, pharmacology, and transport kinetics of the human serotonin transporter. J Biol Chem 293: 3510–3523

Li J, Tajkhorshid E (2009) Ion-releasing state of a secondary membrane transporter. Biophys J 97: L29–31

Li J, Zhao Z, Tajkhorshid E (2019) Locking Two Rigid-body Bundles in an Outward-Facing Conformation: The Ion-coupling Mechanism in a LeuT-fold Transporter. Sci Rep 9: 19479

Liu S, Chang S, Han B, Xu L, Zhang M, Zhao C, Yang W, Wang F, Li J, Delpire E et al (2019) Cryo-EM structures of the human cation-chloride cotransporter KCC1. Science 366: 505–508

Lytle C, McManus TJ, Haas M (1998) A model of Na-K-2Cl cotransport based on ordered ion binding and glide symmetry. Am J Physiol 274: C299–309

Mackenzie B, Ujwal ML, Chang MH, Romero MF, Hediger MA (2006) Divalent metal-ion transporter DMT1 mediates both H+-coupled Fe2+ transport and uncoupled fluxes. Pflugers Arch 451: 544–558

Malinauskaite L, Quick M, Reinhard L, Lyons JA, Yano H, Javitch JA, Nissen P (2014) A mechanism for intracellular release of Na+ by neurotransmitter/sodium symporters. Nat Struct Mol Biol 21: 1006–1012

Malinauskaite L, Said S, Sahin C, Grouleff J, Shahsavar A, Bjerregaard H, Noer P, Severinsen K, Boesen T, Schiott B et al (2016) A conserved leucine occupies the empty substrate site of LeuT in the Na(+)-free return state. Nat Commun 7: 11673

Markadieu N, Delpire E (2014) Physiology and pathophysiology of SLC12A1/2 transporters. Pflugers Arch 466: 91–105

Monette MY, Forbush B (2012) Regulatory activation is accompanied by movement in the C terminus of the Na-K-Cl cotransporter (NKCC1). J Biol Chem 287: 2210–2220

Nakane T, Kimanius D, Lindahl E, Scheres SH (2018) Characterisation of molecular motions in cryo-EM single-particle data by multi-body refinement in RELION. Elife 7

Payne JA (2012) Molecular operation of the cation chloride cotransporters: ion binding and inhibitor interaction. Curr Top Membr 70: 215–237

Penmatsa A, Wang KH, Gouaux E (2013) X-ray structure of dopamine transporter elucidates antidepressant mechanism. Nature 503: 85–90

Perez C, Koshy C, Yildiz O, Ziegler C (2012) Alternating-access mechanism in conformationally asymmetric trimers of the betaine transporter BetP. Nature 490: 126–130

Punjani A, Rubinstein JL, Fleet DJ, Brubaker MA (2017) cryoSPARC: algorithms for rapid unsupervised cryo-EM structure determination. Nat Methods 14: 290–296

Race JE, Makhlouf FN, Logue PJ, Wilson FH, Dunham PB, Holtzman EJ (1999) Molecular cloning and functional characterization of KCC3, a new K-Cl cotransporter. Am J Physiol 277: C1210–1219

Reid MS, Kern DM, Brohawn SG (2020) Cryo-EM structure of the potassium-chloride cotransporter KCC4 in lipid nanodiscs. Elife 9

Rohou A, Grigorieff N (2015) CTFFIND4: Fast and accurate defocus estimation from electron micrographs. J Struct Biol 192: 216–221

Russell JM (2000) Sodium-potassium-chloride cotransport. Physiol Rev 80: 211–276

Scheres SH (2012) RELION: implementation of a Bayesian approach to cryo-EM structure determination. J Struct Biol 180: 519–530

Sen N, Shi L, Beuming T, Weinstein H, Javitch JA (2005) A pincer-like configuration of TM2 in the human dopamine transporter is responsible for indirect effects on cocaine binding. Neuropharmacology 49: 780–790

Shaffer PL, Goehring A, Shankaranarayanan A, Gouaux E (2009) Structure and mechanism of a Na+-independent amino acid transporter. Science 325: 1010–1014

Shahsavar A, Stohler P, Bourenkov G, Zimmermann I, Siegrist M, Guba W, Pinard E, Sinning S, Seeger MA, Schneider TR et al (2021) Structural insights into the inhibition of glycine reuptake. Nature 591: 677–681

Shaikh SA, Tajkhorshid E (2010) Modeling and dynamics of the inward-facing state of a Na+/Cl-dependent neurotransmitter transporter homologue. PLoS Comput Biol 6

Simon DB, Karet FE, Hamdan JM, DiPietro A, Sanjad SA, Lifton RP (1996a) Bartter’s syndrome, hypokalaemic alkalosis with hypercalciuria, is caused by mutations in the Na-K-2Cl cotransporter NKCC2. Nat Genet 13: 183–188

Simon DB, Nelson-Williams C, Bia MJ, Ellison D, Karet FE, Molina AM, Vaara I, Iwata F, Cushner HM, Koolen M et al (1996b) Gitelman’s variant of Bartter’s syndrome, inherited hypokalaemic alkalosis, is caused by mutations in the thiazide-sensitive Na-Cl cotransporter. Nat Genet 12: 24–30

Somasekharan S, Tanis J, Forbush B (2012) Loop diuretic and ion-binding residues revealed by scanning mutagenesis of transmembrane helix 3 (TM3) of Na-K-Cl cotransporter (NKCC1). J Biol Chem 287: 17308–17317

Sucic S, Paczkowski FA, Runkel F, Bonisch H, Bryan-Lluka LJ (2002) Functional significance of a highly conserved glutamate residue of the human noradrenaline transporter. J Neurochem 81: 344–354

Tavoulari S, Margheritis E, Nagarajan A, DeWitt DC, Zhang YW, Rosado E, Ravera S, Rhoades E, Forrest LR, Rudnick G (2016) Two Na+ Sites Control Conformational Change in a Neurotransmitter Transporter Homolog. J Biol Chem 291: 1456–1471

Turner RJ, George JN (1988) Ionic dependence of bumetanide binding to the rabbit parotid Na/K/Cl cotransporter. J Membr Biol 102: 71–77

Wahlgren WY, Dunevall E, North RA, Paz A, Scalise M, Bisignano P, Bengtsson-Palme J, Goyal P, Claesson E, Caing-Carlsson R et al (2018) Substrate-bound outward-open structure of a Na(+)-coupled sialic acid symporter reveals a new Na(+) site. Nat Commun 9: 1753

Warmuth S, Zimmermann I, Dutzler R (2009) X-ray structure of the C-terminal domain of a prokaryotic cation-chloride cotransporter. Structure 17: 538–546

Watanabe A, Choe S, Chaptal V, Rosenberg JM, Wright EM, Grabe M, Abramson J (2010) The mechanism of sodium and substrate release from the binding pocket of vSGLT. Nature 468: 988–991

Weyand S, Shimamura T, Yajima S, Suzuki S, Mirza O, Krusong K, Carpenter EP, Rutherford NG, Hadden JM, O’Reilly J et al (2008) Structure and molecular mechanism of a nucleobase-cation-symport-1 family transporter. Science 322: 709–713

Williams CJ, Headd JJ, Moriarty NW, Prisant MG, Videau LL, Deis LN, Verma V, Keedy DA, Hintze BJ, Chen VB et al (2018) MolProbity: More and better reference data for improved all-atom structure validation. Protein Sci 27: 293–315

Xie Y, Chang S, Zhao C, Wang F, Liu S, Wang J, Delpire E, Ye S, Guo J (2020) Structures and an activation mechanism of human potassium-chloride cotransporters. Sci Adv 6

Yamashita A, Singh SK, Kawate T, Jin Y, Gouaux E (2005) Crystal structure of a bacterial homologue of Na+/Cl--dependent neurotransmitter transporters. Nature 437: 215–223

Yan R, Zhang Y, Li Y, Xia L, Guo Y, Zhou Q (2020) Structural basis for the recognition of SARS-CoV-2 by full-length human ACE2. Science 367: 1444–1448

Yang X, Wang Q, Cao E (2020) Structure of the human cation-chloride cotransporter NKCC1 determined by single-particle electron cryo-microscopy. Nat Commun 11: 1016

Zhang S, Zhou J, Zhang Y, Liu T, Friedel P, Zhuo W, Somasekharan S, Roy K, Zhang L, Liu Y et al (2021) The structural basis of function and regulation of neuronal cotransporters NKCC1 and KCC2. Commun Biol 4: 226

Zhang YW, Tavoulari S, Sinning S, Aleksandrova AA, Forrest LR, Rudnick G (2018) Structural elements required for coupling ion and substrate transport in the neurotransmitter transporter homolog LeuT. Proc Natl Acad Sci U S A 115: E8854–E8862

Zhao Y, Terry D, Shi L, Weinstein H, Blanchard SC, Javitch JA (2010) Single-molecule dynamics of gating in a neurotransmitter transporter homologue. Nature 465: 188–193

Zimanyi CM, Guo M, Mahmood A, Hendrickson WA, Hirsh D, Cheung J (2020) Structure of the Regulatory Cytosolic Domain of a Eukaryotic Potassium-Chloride Cotransporter. Structure

